# Defined Mars Media (DMM), a chemically defined simulant of the soluble macro- and micro- nutrients in Mars regolith for use in biological research

**DOI:** 10.64898/2026.04.24.719001

**Authors:** Harley Greene, Una Nattermann, Devon A. Stork, Fatima R. Martin, Max G. Schubert, Tom Pedersen, Edward Sukarto, Amy Spens, Jordan E. Mancuso, Keren Isaev, Nathan D. Hicks, Jonathan Liu, Rachel Harris, Charles Cockell, Samuel P. Kounaves, Erika A. DeBenedictis

**Author notes:** authors contributed equally to this work (Harley Greene, Una Nattermann).

## Abstract

Mars’ harsh yet workable surface conditions, such as manageable temperatures, availability of solar energy, and *in situ* resources like water ice, carbon dioxide, and mineral-rich regolith, make it a compelling target for supporting life beyond Earth. However, existing experiments testing chemical habitability in Mars conditions generally rely on leachates of physical regolith simulants, which vary in composition across simulant types, leaching conditions, and production batches. We introduce a defined Mars media (DMM) that accurately simulates the biologically relevant nutrients (nitrogen, phosphorus, and sulfur) and stressors (perchlorates, heavy metals) in Martian regolith when it is leached in water at neutral pH. We formulated DMM by combining direct rover and lander measurements from Mars with laboratory measurements of regolith simulant leachates. We validate DMM from a 1x to 20x concentrate, equivalent to 40 g/L to 800 g/L of leached regolith. Using DMM with acetate as a Mars atmosphere-derived carbon source, we grew eight bacteria, demonstrating that organisms can source all essential nutrients from Martian resources. We also demonstrate that microbial growth in DMM is robust to uncertainties in Martian regolith composition: sensitivity experiments can identify limiting trace element nutrients and toxins in DMM, and show that bacterial growth is maintained across at least an order of magnitude variation in their concentrations. This is the first defined Mars regolith media recipe containing both macro- and micro-nutrients, and designed specifically for biological experimentation. By shifting from variable leachate-based approaches to a defined aqueous analog, we enable controlled hypothesis testing of microbial survival, growth, and function in a Martian chemical environment. DMM will enable further research on astrobiology, biological *in situ* resource utilization, large-scale soil remediation, and terraforming.

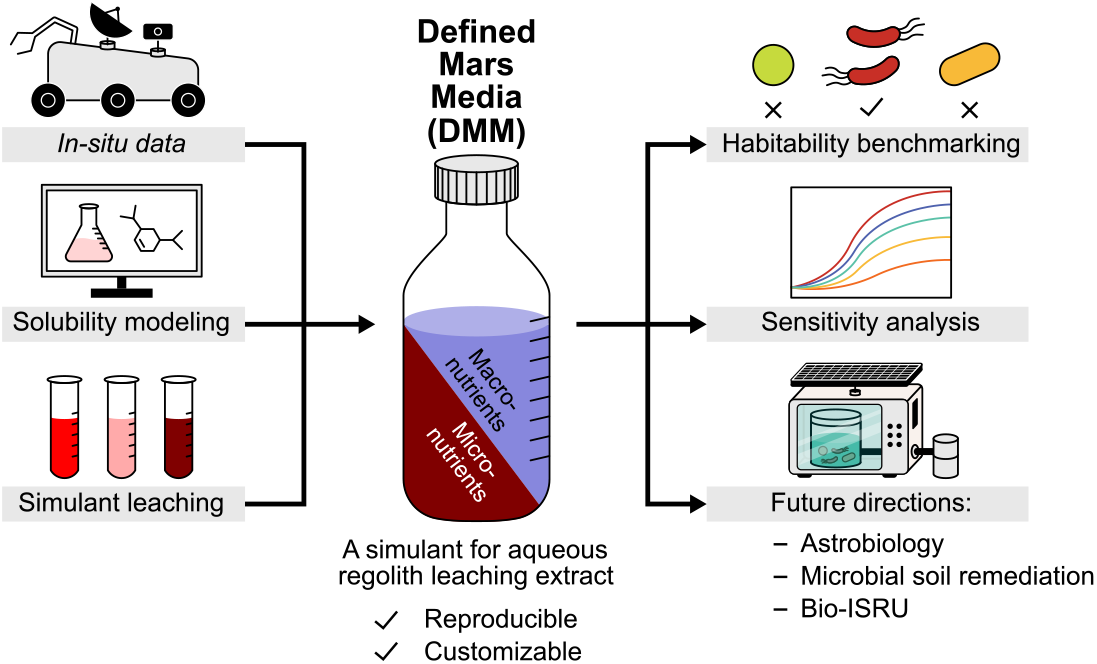

## INTRODUCTION

Determining chemical habitability in *in situ* Martian resources is a central question in modern space science, aiming to understand if the existing chemicals on Mars can support microbial life. Traditionally, this question has been investigated through the lens of astrobiology and the search for extant life on Mars, which remains the highest-priority science objective for the first human mission to Mars^1^.However, understanding microbial chemical habitability on Mars is increasingly discussed through the lens of a second context: deliberate use of biological engineering for manufacturing. As exploration architectures evolve from robotic precursors^2^ toward sustained human presence^3^, the ability to source materials locally becomes increasingly critical to enable Earth-independence and mission resilience^4,5^. *In situ* Resource Utilization (ISRU) could create useful materials from local resources without costly and slow resupply from Earth^4,6–8^. Biomanufacturing is a particularly promising ISRU technology with the potential to produce a wide variety of useful materials at scale^4^ and create habitable environments for humans^6–8^. In both cases, whether or not microbes can rely on the available chemical resources to proliferate is of central importance to the search-for, and use-of, life on Mars, and is a key area of research at Pioneer Labs (pioneer-labs.org).

A major outstanding question is whether there exists a sufficient *in situ* feedstock on Mars to fully support life with minimal infrastructure. The atomic building blocks for life (i.e. CHNOPS) are present in the Martian atmosphere, regolith, and ice at massive scale^9–12 .^However, elemental presence alone does not guarantee biological habitability^9–13^. Physical and chemical factors such as oxidizing species, perchlorates and other potential stressors in regolith, extreme salinity and pH regimes, limited liquid water availability, and intense radiation may severely constrain microbial survival and metabolism even with plentiful raw materials^14–19.^A microbe capable of growing on Mars must both extract sufficient nutrients and survive stressors, or be engineered to do so, until human involvement allows for abiotic material conversion (ex. fertilizer production for use in agriculture).

Martian regolith is a particularly promising feedstock component as a source of inorganic nutrients such as nitrogen, phosphorus, sulfur, and trace metals, while the atmosphere is the primary source of carbon via carbon dioxide. The Martian atmosphere does contain gaseous nitrogen (2.85% of the atmosphere^20^), but no organism has been identified that is able to fix N_2_ at the current partial pressure on Mars’ surface (pN_2_ < 1 mbar)^21,22^ . Deconvoluting Martian regolith’s chemical makeup to understand the biologically accessible and relevant components is essential for understanding chemical habitability and bio-ISRU efforts. There has only been one aqueous chemical analysis of regolith to date, by the Phoenix Lander’s Wet Chemistry Laboratory (WCL), which provided critical insights into bio-available soluble components, like the discovery of perchlorate^14,23^ There are no future plans for similar measurements, beyond potential Mars Sample Return, although near-term funding and implementation pathways remain uncertain^24^. Numerous Mars regolith simulants, or analogs, have been developed over the past decades to mimic regolith’s physical characteristics and mineralogical composition^25^. Some simulants have been used in microbiology-focused studies, commonly via aqueous extraction or leaching, to assess habitability in nutrients sourced from the regolith simulant-derived solutions and tolerance to soluble toxic species **(Table 1)**. Collectively, this body of work has helped establish proof-of-concept that microbial activity is possible in Mars-relevant chemical contexts across microorganisms and regolith simulants. However, the choice of simulant and extraction methodology vary considerably across studies, with both factors significantly influencing the recoverable soluble nutrient concentrations (Appendix I). Furthermore, these mineral simulants were developed for non-biological purposes, such as lander and rover development and abiotic ISRU (ex. water extraction). For example, all of the existing commercial simulants lack perchlorate salts; and thus, astrobiology researchers typically add perchlorate salts to the mineral simulant leachates^26,27^. While these mineral simulants mimic bulk solid-phase characteristics, it is essential to simulate defined concentrations of “bioavailable” resources, those components of the regolith that dissolve in solution and can be accessed by microbes, for biological research. Therefore, it stands to reason that *in* situ data detailing the biologically relevant components of regolith should have higher consideration when investigating habitability in regolith-derived chemicals.

**Table 1.**
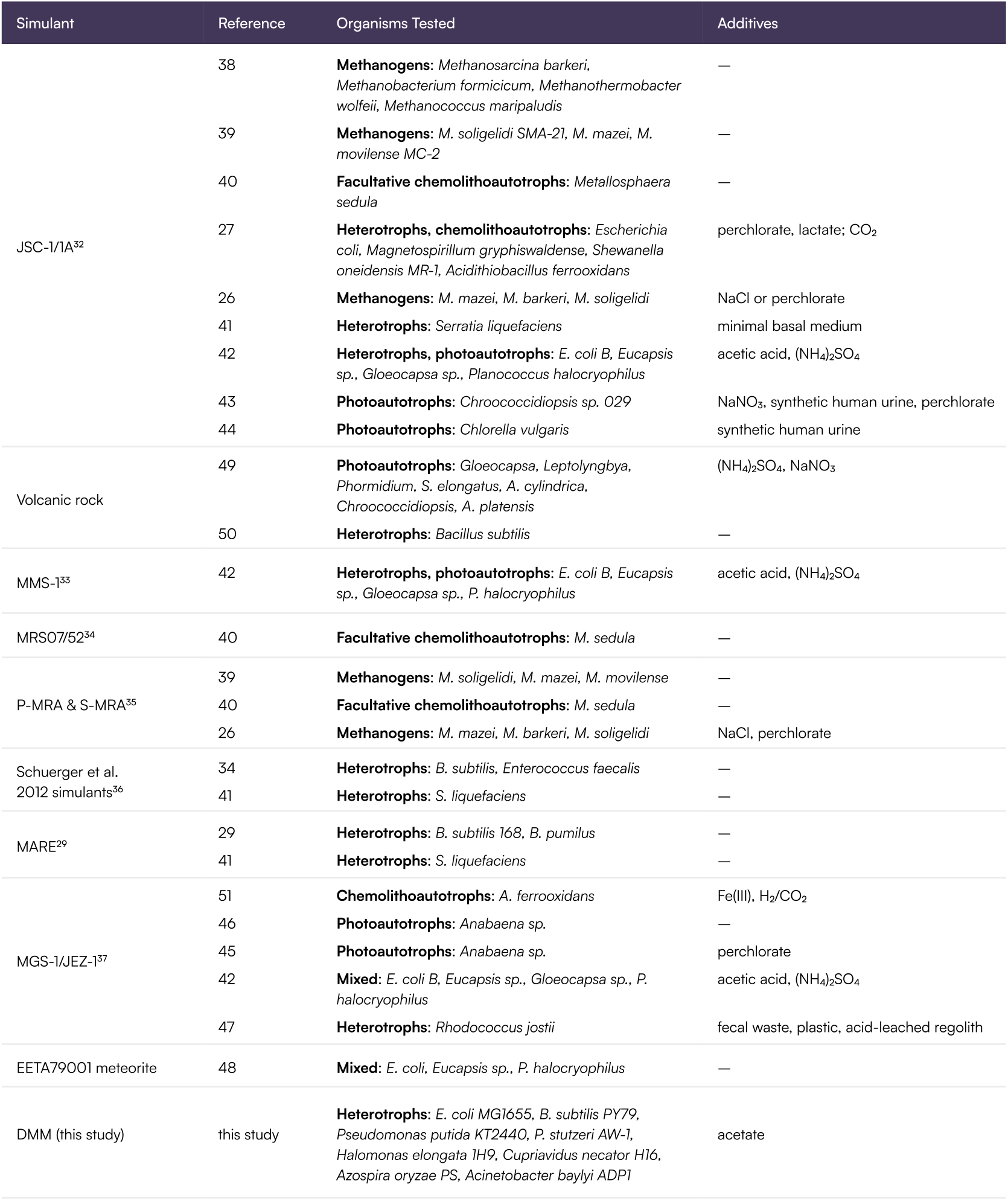
Summary of relevant microbial chemical habitability studies using existing Mars regolith simulants, including the organisms tested and any added supplements permitting growth^26,27,29,32–51^.

For high-throughput screening, adaptive evolution, and synthetic biology experiments, it is essential to model all chemical species with high fidelity in a reproducible and controllable manner. This underscores the pressing need for a chemically defined, standardized soluble Mars regolith medium for microbiology^28^. Routinely, microbiologists design media to reflect specific environments, such as artificial sea water, an enrichment condition for bioprospecting, or ideal cultivation conditions for a particular strain (ex. https://mediadive.dsmz.de/). Defined chemical media exist to provide reproducible nutrient backgrounds in which to conduct growth and engineering studies. Previously, only one chemically defined soluble Mars regolith analog recipe has been published for microbiological experimentation: a Mars Simulant that matches Phoenix WCL chemistry^29,30^. However, Mars Simulant is an incomplete nutrient source and is out of date to current research, as it contains ammonium and lacks nitrates and phosphates^9,11,31^. To approach formulating a chemically defined Mars regolith media designed for microbiology and chemical habitability work, we consulted existing Mars chemical data and modeling literature.

Here, we present Defined Mars Media (DMM), a reproducible mimic of aqueous Martian regolith chemistry, designed explicitly for microbial research using Martian resources (Box 1). Our goal is a medium grounded in Mars mission data and modeling literature that captures relevant nutrient availability and stressor chemistry. When *in situ* aqueous data is not available, we prioritized maintaining Mars relevancy by using available or in-house generated dissolution data on regolith analogs and we describe incorporating supplemental chemicals depending on the use case. Establishing a standard medium recipe allows for modification or customization based on new data or specific research needs. We demonstrate DMM’s applicability for astrobiology research, such as studying microbial chemical habitability and identifying growth limiting nutrients or stressors. The base medium can be systematically adjusted to represent different plausible Martian locales, water activities, or mission scenarios, supporting both mechanistic studies and iterative strain engineering under well-specified conditions. DMM will enable reproducible experimentation across labs, high throughput chemical habitability studies, and supply a modifiable media for evolution and engineering efforts.

### BOX 1.

How to prepare defined Mars media (DMM)

**Preparation Of DMM**

DMM is prepared in four steps:

1. Prepare a macronutrient solution from stock solutions
2. Prepare a concentrated micronutrient solution and add to diluted macronutrients
3. Add in supplements as needed
4. Add water to desired volume and concentration

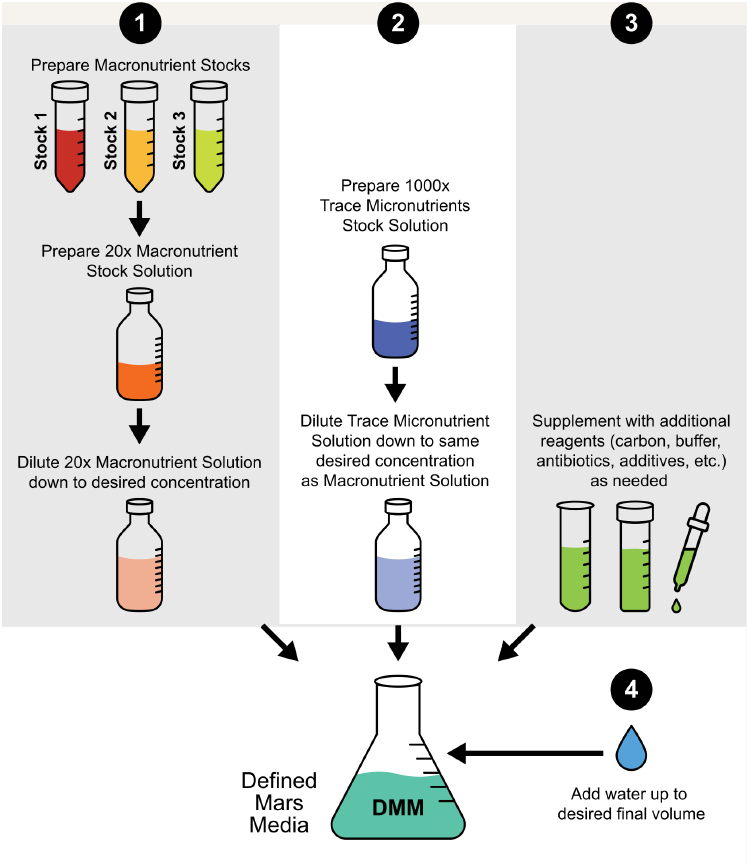

**Preparation of a 20x Macronutrient Solution**

First, prepare the three Macronutrient Stock Solutions (500x Bulk Salts, 1000x Perchlorate Salts, 1000x Phosphate Salts) according to the concentration in the table below. Dilute the three Stock Solutions together in sterile deionized water to the desired DMM concentration (examples for 1x and 20x are shown below). Add Stock 2 and Stock 3 slowly with constant mixing to reduce precipitation of calcium sulfate and magnesium phosphate. At ≥20x DMM, precipitation may be unavoidable. Sterilize with a 0.2 μm filter and store for up to six months.

| Stock Number | Ingredients | Volume For 1 L Of 1x DMM | Volume For 1 L Of 20x DMM |
| --- | --- | --- | --- |
| 1 | 23.08 g/L NaCl<br>25.28 g/L KNO<sub>3</sub><br>21.25 g/L NaNO<sub>3</sub><br>164.48 g/L Na<sub>2</sub>SO<sub>4</sub> · 10H<sub>2</sub>O<br>367.13 g/L MgSO<sub>4</sub> · 7H<sub>2</sub>O | 2 mL | 40 mL |
| 2 | 91.11 g/L Mg(ClO<sub>4</sub>)<sub>2</sub> · 6H<sub>2</sub>O<br>362.92 g/L Ca(ClO<sub>4</sub>)<sub>2</sub> · 4H<sub>2</sub>O | 1 mL | 20 mL |
| 3 | 18.23 g/L Na<sub>2</sub>HPO<sub>4</sub> · 7H<sub>2</sub>O<br>4.42 g/L NaH<sub>2</sub>PO<sub>4</sub> · H<sub>2</sub>O | 1 mL | 20 mL |
|  | Sterile water | up to 1 L | up to 1 L |

**Preparation of a 1000x DMM Trace Micronutrient Stock Solution**

Prepare stock solutions of individual trace elements as described in the “Stock Solution” column in the table below. To prepare the 1000x stock of trace micronutrients, start with 850 mL of DI water, add EDTA & HCl to obtain pH 6. Add solid FeSO_4_ · 7H_2_O and stir until dissolved. Add the remaining components volumetrically with constant mixing and bring the solution to a 1 L final volume. Sterilize with a 0.2 μm filter, wrap in foil to prevent photodegradation, and store at 4°C until use.

| Chemical | Stock Solution | Quantity For 1 L Of 1000x Stock | Molar Concentration In Final 1000x Stock (mM) |
| --- | --- | --- | --- |
| EDTA, pH 8 | 500 mM | 33 mL | 16.5 |
| HCl |  | to pH 6 |  |
| FeSO<sub>4</sub> · 7H<sub>2</sub>O | — | 2 g | 7.1940 |
| AlK(SO<sub>4</sub>)<sub>2</sub> · 12H<sub>2</sub>O | 100 g/L | 40 mL | 8.4319 |
| SrCl<sub>2</sub> · 6H<sub>2</sub>O | 200 g/L | 10 mL | 7.5013 |
| ZnSO<sub>4</sub> · 7H<sub>2</sub>O | 300 g/L | 1 mL | 1.0433 |
| H<sub>3</sub>BO<sub>3</sub> | 30 g/L | 2 mL | 0.91 |
| LiCl | 10 g/L | 1 mL | 0.2359 |
| BaCl<sub>2</sub> · 2H<sub>2</sub>O | 20 g/L | 1 mL | 0.0819 |
| Na<sub>2</sub>MoO<sub>4</sub> · 2H<sub>2</sub>O | 10 g/L | 1 mL | 0.0413 |
| CrCl<sub>3</sub> · 6H<sub>2</sub>O | 10 g/L | 1 mL | 0.0375 |
| CuCl<sub>2</sub> · 2H<sub>2</sub>O | 6 g/L | 1 mL | 0.0352 |
| NiCl<sub>2</sub> · 6H<sub>2</sub>O | 8 g/L | 1 mL | 0.0337 |
| VO<sub>2</sub>SO<sub>4</sub> · nH<sub>2</sub>O | 3 g/L | 1 mL | 0.0184 |
| MnCl<sub>2</sub> · 4H<sub>2</sub>O | 0.3 g/L | 10 mL | 0.0152 |
| Na<sub>2</sub>SeO<sub>3</sub> | 2 g/L | 1 mL | 0.0116 |
| Na<sub>2</sub>WO<sub>4</sub> · 2H<sub>2</sub>O | 2 g/L | 1 mL | 0.0061 |
| CoCl<sub>2</sub> · 6H<sub>2</sub>O | 0.5 g/L | 2 mL | 0.0042 |
| Zr(SO<sub>4</sub>)<sub>2</sub> · 4H<sub>2</sub>O | 1 g/L | 1 mL | 0.0028 |
| AgNO<sub>3</sub> | 3 g/L | 0.01 mL | 0.0002 |

**Preparation of complete working media**

Examples are shown in the table below for preparing a 1x DMM and 10x DMM from the 20x Macronutrient Solution, 1000x Trace Micronutrient Stock Solution, and concentrated carbon and buffer sources. Add carbon and buffer slowly with constant mixing to reduce precipitation.

See Supplementary Table 1 for a list of manufacturers and catalog numbers for the reagents we purchased to make DMM.

| Stock | Volume For 1 L Of 1x DMM, 1% K-Acetate, 50 mM TES | Volume For 1 L Of 10x DMM, 1% K-Acetate, 50 mM TES |
| --- | --- | --- |
| 20x DMM Macronutrient Stock Solution | 50 mL | 500 mL |
| 1000x Trace Micronutrient Stock Solution | 1 mL | 10 mL |
| 20% Potassium acetate | 50 mL | 50 mL |
| 1 M TES, pH 8.0 | 50 mL | 50 mL |
| Sterile water | up to 1 L | up to 1 L |

## DEFINED MARS MEDIA (DMM)

Chemically defined media for biology is composed of pure organic and inorganic components, covering all biological necessities like salts, carbon sources, buffering agents, vitamins, and trace elements. Our approach to formulating a defined aqueous simulant follows a similar framework, but rather than optimizing the medium for bacterial growth, we aim to reproduce the soluble fraction expected from aqueous extraction of Mars regolith.

Here we present DMM, an aqueous Mars regolith simulant modeling the concentration of major and minor soluble species. The baseline concentration, which we refer to as lx DMM, simulates the soluble chemistry of 40 g of Martian regolith added to 1 x L of water, equivalent to the *in situ* measurement conditions by the Phoenix WCL^14,23^. To create our recipe, we break DMM into two parts: a macronutrient portion describing the available major soluble species and a micronutrient portion describing the soluble trace elements in regolith. Because DMM is essentially a salt mixture, it can be modified to probe individual chemical species depending on the intended use case or organism of interest. We also describe how to customize DMM by supplementing it with ISRU-derived carbon sources or buffering agents. making it compatible with heterotrophic bacteria chemical habitability experiments.

### Formulating DMM Macronutrient Profile

We define the macronutrients as the major soluble species above 1 ppm found when mixing Mars regolith with water. We formulated the DMM macronutrient profile around Phoenix Lander WCL data and associated modeling. Curiosity Rover Sample Analysis at Mars (SAM) mineralogical data, and literature on leachates of Phoenix Lander site regolith mimics. The Phoenix WCL consisted of sensors to measure the aqueous chemistry of the sampled regolith at 5-10°C and ∼8.4 mbar and was designed to detect cations (Na^+^, K^+^, Mg^2+^, Ca^2+^, NH4^+^), anions (Cl^−^, Br^−^,I^−^, NO_3_^−^, SO_4_ ^2-^). specific heavy metals (Cu^2+^, Cd^2+^, Pb^2+^, Fe^213+^, Hg^2+^), and pH^52^. Other sensors. like electrodes for Ba^2+^ and Li^+^, were included for calibration purposes. We used averaged data reported from the Phoenix WCL on soluble ionic species that were above detection limits. equilibrium model analysis of that data. and later-on reports of Phoenix site aqueous chemistry to create the basis for our macronutrient formulation^14,23,30,48,53–55^ **(Table 2)**. In the WCL tests. sampled regolith was mixed with water at a 1:25 ratio of regolith to water, equivalent to 40 g regolith / L water, and we therefore use the same ratio as our standard. During Phoenix WCL. the Hofmeister electrode for nitrate detection revealed the presence of perchlorate ions^14^ Critically, perchlorates are not included in the commercial Mars regolith simulants. but are considered to be globally distributed on Mars with some regional variation^14,19,56–60^. Therefore, we use Phoenix WCL values for the perchlorates in our defined media. Our leaching studies of commercial mineral regolith simulants indicate that Earth pressure and room temperature (∼20-25°C) result in similar soluble ions to the Phoenix WCL reports, except perchlorate which is not included in physical simulants, and we explore leaching variables like pH, concentration of added regolith, and leaching time **(Appendix I)**.

**Table 2.** Composition of the major soluble ions (macronutrients) in our defined Mars media.

| Ion | Chosen Concentration (mM) | Lower Bound (mM) | Average (mM) | Upper Bound (mM) | References Used For Formulation |
| --- | --- | --- | --- | --- | --- |
| Na<sup>+</sup> | 3.50 | 0.99 | 1.49 | 3.52 | 14, 23, 30, 48, 53, 54, 55 |
| K<sup>+</sup> | 0.50 | 0.17 | 0.34 | 0.50 | 14, 23, 30, 48, 53, 54, 55 |
| Ca<sup>2+</sup> | 1.17 | 0.09 | 0.52 | 1.17 | 9, 14, 23, 30, 48, 53, 54, 55 |
| Mg<sup>2+</sup> | 3.25 | 1.31 | 3.63 | 6.40 | 14, 23, 30, 48, 53, 54, 55 |
| Cl<sup>-</sup> | 0.79 | 0.24 | 0.45 | 0.79 | 14, 23, 30, 48, 53, 54 |
| ClO<sub>4</sub><sup>-</sup> | 2.89 | 2.10 | 2.42 | 2.89 | 14, 23, 30, 48, 53, 54 |
| SO<sub>4</sub><sup>2-</sup> | 4.00 | 4.79 | 4.80 | 5.90 | 30, 53, 54, 55 |
| NO<sub>3</sub><sup>-</sup> | 1.00 | 0.64 | 0.88 | 1.00 | 30, 48, 55 |
| PO<sub>4</sub><sup>3-</sup> | 0.10 | 0.096 | 0.10 | 0.10 | 9 |
| pH | 7.70 | 6.52 | 7.50 | 7.74 | 14, 23, 30, 54 |

While carbonate-containing minerals have been reported by Curiosity SAM and orbital data, bicarbonate (HCO_3_^-^) shows up in some, but not all. equilibrium models of the Phoenix WCL data^12,53,54,61,62^. Additionally, studies where carbonate minerals were used to simulate Mars regolith do not report soluble bicarbonate in their experimental characterization^30,55^, likely because the Phoenix WCL experiment was conducted at Mars temperatures and CO2 partial pressures. while equivalent studies in Earth conditions have lower bicarbonate solubility. We therefore chose not to include *in situ* bicarbonates in the DMM formulation.

As mentioned, the Phoenix WCL included a Hofmeister electrode, but the experiment was not able to produce quantitative nitrate measurements due to interference with the high concentrations of perchlorates^14,23^. However, later Curiosity SAM data shows that there are fixed nitrogen species, including nitrates, in the regolith, and these would likely be highly soluble under aqueous conditions^11,56^. Due to the presence of nitrate containing minerals on Mars and the measurement of soluble nitrates in synthetic leachates, we decided to add nitrate into our DMM formulation^30,48,55^. Ammonium was reported in initial WCL data^14^.but was measured at less than the limit of detection for all three soil samples^23^.

Ammonium was expected as a contaminant from the lander’s descent thruster fuel, not an indigenous source of fixed nitrogen^31^. Therefore, we did not include ammonium in the DMM formulation. Previous Mars simulants notably do not model the presence of soluble phosphates. This is important, since regolith is the only possible in *situ* phosphorus source for living organisms. We know there are abundant phosphate minerals on Mars from alpha particle X-ray spectrometry data^63,64^. More recent measurements by the Perseverance rover identified iron-phosphate minerals associated with organic-rich aqueous-altered sediments in Jezero Crater^65^. As of yet, there is no *in situ* data on mineral versus bioavailable phosphorus. However, researchers performed dissolution experiments on representative apatite minerals and measured soluble phosphate^9^. We were encouraged by these studies and added soluble phosphorus, as phosphate, into the DMM formulation.

Taken together, we ended up with an average, lower, and upper bound for the macronutrient species including small amounts of soluble nitrate and phosphate in our DMM formulation **(Table 2)**. We chose the final ion concentrations based on making a salt recipe to achieve an ionic formulation while ensuring a charge balance of pH 7.7, as reported by Phoenix WCL^14,23^**(Box 1**).

### In-House Leaching Of Commercial Simulants And Elemental Analysis To Produce A Micronutrient Profile

Beyond macronutrients, organisms depend on trace metals for several core biological functions. For example, iron is an essential metal cofactor for all respiring organisms and manganese is important for functions like metabolism and oxidative stress resistance, but both are only required in trace amounts (less than 1 ppm)^66,67^. Other trace elements may be necessary for nonessential yet Mars-relevant functions: for example, anaerobic prokaryotes utilizing tungsten to survive in high salinity brines^68-69.^ Several elements commonly defined as nonessential can still be biologically relevant in certain conditions, such as vanadium as an alternative nitrogenase cofactor^70,71^.

In lieu of any existing *in situ* aqueous chemistry data on trace elements in regolith, we performed in-house leaching of commercial Mars regolith simulants and sent the extracts for elemental analysis using inductively coupled plasma mass spectrometry (ICP-MS) to approximate aqueous trace elements. A critical caveat applies: commercial simulants are designed to replicate the bulk mineralogy and physical properties of Martian regolith, not its trace element chemistry. Their trace profiles are shaped by the source geology of their terrestrial parent material, which may or may not reflect Martian conditions. Whether Earth rocks are valid geochemical proxies for Mars rocks is arguably the central open question in planetary science, with evidence on both sides: similar silicate and oxide mineral phases exist in both planets’ crusts, but different histories of aqueous alteration, biological cycling, and mantle evolution could shift trace element partitioning in ways that remain unresolved without *in situ* measurement^72–75^. Mars meteorite analyses provide a closer geochemical proxy but still represent igneous lithologies rather than surface granular media^76–78^. Therefore, we use these data to identify which elements could reasonably be mobilized from silicate and oxide mineral phases broadly common to both planets, while flagging that absolute concentrations remain unconstrained in the absence of *in situ* soluble trace element measurements. This reflects our modular formulation approach: the previously described macronutrient portion reflecting the species and concentrations documented by *in situ* instrument data, supplemented by an optional, yet still Mars-relevant, trace element solution at concentrations informed by simulant leaching, which can be included, excluded, or modified depending on the experiment in question.

To formulate this trace element solution, we prepared extracts from three different commercially available regolith simulants in neutral pH water at 40 g/L regolith (MGS-1, JEZ-7, MMS-2; see **Methods** and **Supplementary Table 2**). We considered three categories of soluble trace elements: biologically relevant elements (Mn, Cu, W, B, Fe, Zn, Ni, Mo, Co, Se), elements included in oxides that make up simulants (ex. Si, Ti, Al), and other Mars-relevant elements present at high concentrations compared to Earth (ex. Cr, Zr, Sr)^76,79,80^. In formulating, we excluded elements that were not consistently measured (i.e. not measured in all three simulant extracts) from the final list of micronutrients, such as Sn, Te, Be, and U. Both Martian regolith and physical simulants contain high amounts of silicon and titanium oxides, but neither metal was detected in neutral leaching conditions, indicating limited solubility. We used the average concentration from the MGS-1 extracts as the formulation concentration as a proxy for global micronutrient presence, and incorporated the other simulants’ extract data to identify maximum and minimum values. We classify the elements measured here as the micronutrient portion of DMM **(Table 3)**.

**Table 3.**
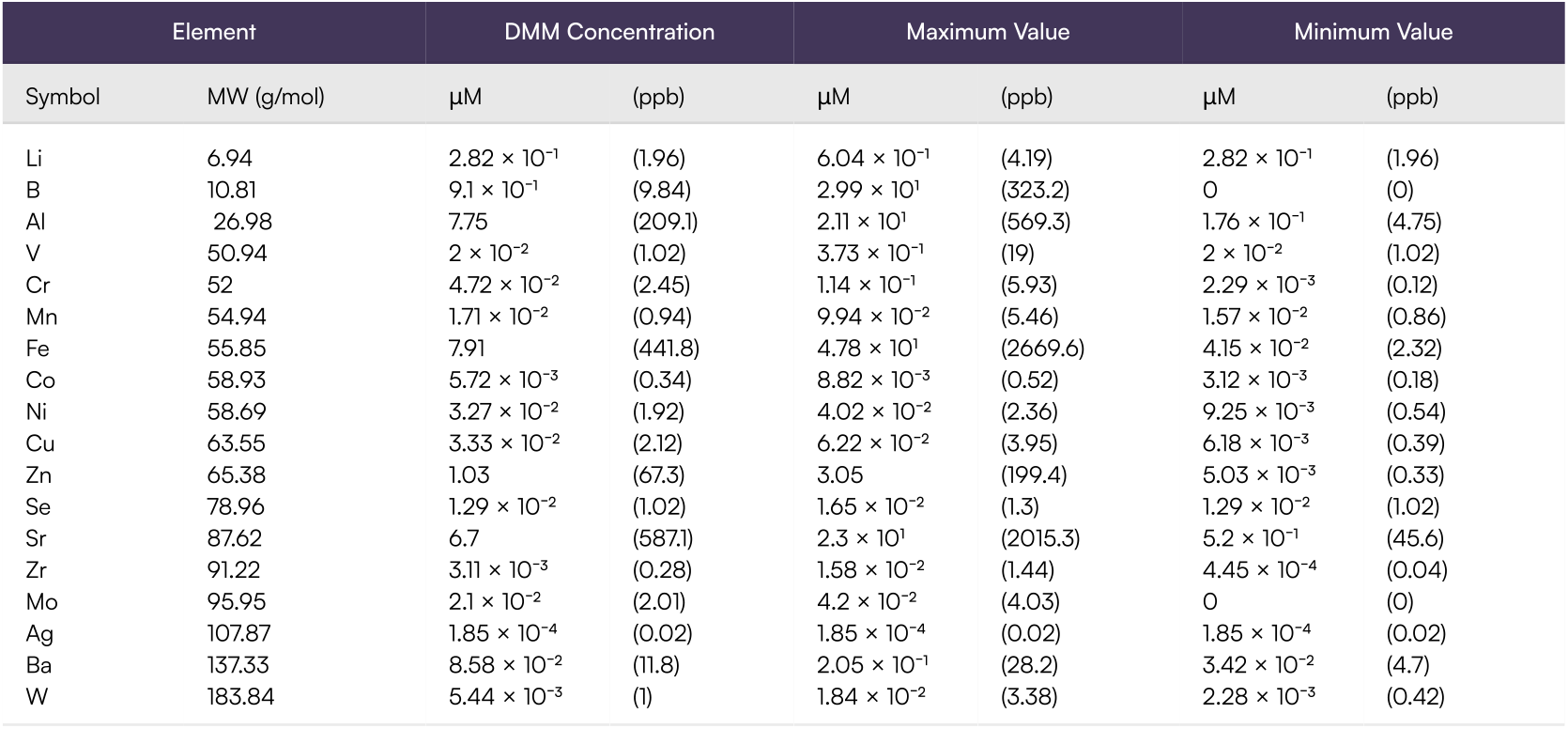
Composition of the soluble micronutrients in DMM, along with the maximum and minimum concentrations measured in leachates of three Mars regolith simulants (MMS-2, MGS-1, and JEZ-1; see **Supplemental Table 2)** at 40 g/L. Values are presented in micromolar and part-per-billion.

We used these data to create a recipe consisting of soluble metal salts to recapitulate a defined micronutrient mix **(Box 1)**. In practice, we make a concentrated micronutrient stock solution and add it into the final media at the desired concentration. There are a few noteworthy aspects of the chosen salts and their impact on the charge state of the soluble elements/ metals. We chose to represent soluble trace iron as Fe(II), as it is more soluble than the oxidized Fe(III), both of which are on Mars, although Fe(III) may be more prominent in the dust and soil on Mars^81–83^. Similarly, we represent soluble trace chromium as Cr(III), the reduced and less toxic ion compared to Cr(VI), for ease of creation and distribution^80,84^. Although Cr(III) is typically two to three times less cytotoxic than Cr(VI), depending on the organism, several bacterial species can flourish in Cr(Vl)-rich environments by reducing it to Cr(lll)^85–88^ . EDTA is added as a chelating agent to ensure metal solubility, similar to its use in standard trace element mixtures, but is not a reliable microbial carbon source.

### Experimental characterization of DMM

Leachates of regolith are, by definition, at the edge of solubility and influenced by non-chemical factors, making formulation and shelf stability of defined leachate media challenging. As with any useful chemically defined media, DMM should be easy to prepare and consistent batch-to-batch. To measure consistency, we made several batches of DMM across different operators and at different times. Batches were sent for ion chromatography (IC) and ICP-MS elemental analysis to measure the observed soluble ionic and elemental concentrations and compared them to the expected formulation concentration. At a lx DMM concentration (i.e. 40 grams of regolith in 1 liter of water), the measured concentrations for total soluble Na, Mg, K, Ca, CI^-^. No_3_^-^, and SO_4_ ^2-^ fall within one standard deviation of the expected formulation **(Figure 1A)**. Note that Na, Mg, K, and Ca are reported as total element concentration as ICP-MS cannot determine charge state. We measured less CIO_4_^-^, than expected and we were not able to accurately quantify soluble phosphate using the available ion chromatography method due to known co-elution issues^149,150^ with the sulfate, phosphate, and perchlorate peaks. However, ion equilibrium modeling supports the presence of the intended perchlorate and phosphate concentrations, as the observed cation balance requires a corresponding anion contribution.

**Figure 1:**
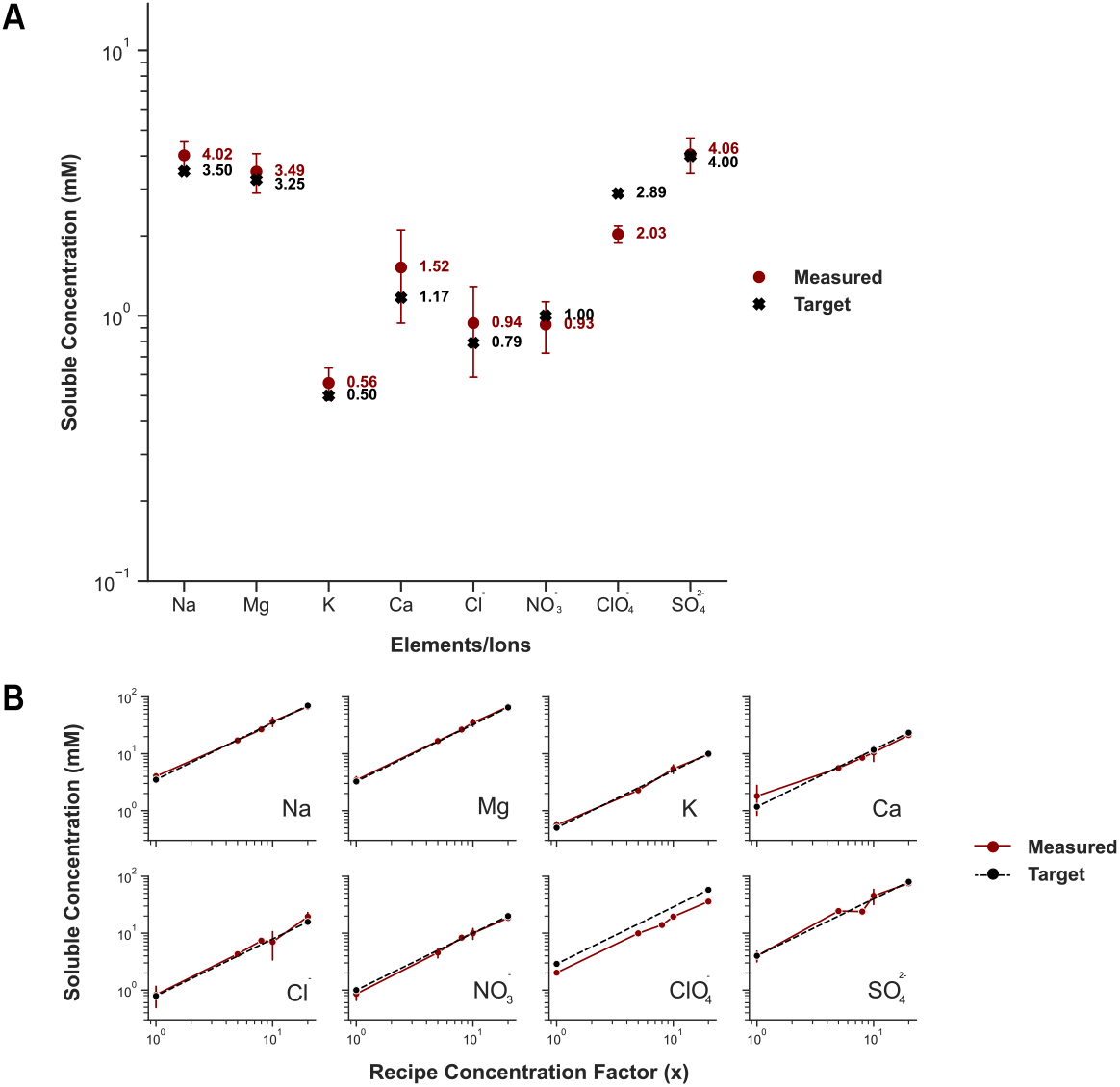
Validation of DMM macronutrients at varying concentrations. A. Comparison of total soluble concentration (red dots, n=7) to target concentration in base formulation (black star) in millimolar for the DMM macronutrients. Phosphate is omitted due to difficulties around large sulfate concentrations impacting IC quantification. B.Comparison of soluble ion concentration measurements (red line) across a range of DMM concentrations to the target concentration in the formulation (black line) for the DMM macronutrients. For the measured data (red line), each dot represents 2-5 measurements, with standard deviation error bars where applicable, and the line connects the averages for each concentration. *For all* Fiqures. *Ion Chromatoqraphv* is *used* for *anion (CI^-^, NO*_*3*_^-^. CIO^-^_4_, PO_4_^3-^, SO_4_ ^2-^) *concentrations and* ICP-MS is *used* for *total element concentrations*.

We sought to prepare DMM at higher concentrations to conduct experiments simulating higher concentration of input regolith, exposing microbes to more bioavailable nutrients and potential soluble stressors. To simulate preparing the extract with higher input concentrations of regolith, DMM batches were prepared at higher concentrations: 5x, 8 x 10 x, and 20x concentration, mimicking the soluble concentrations of 200 g, 320 g, 400 g, and 800 g of regolith in 1 L of water respectively. Higher concentrations of DMM are prepared by multiplying the baseline formulation by the desired concentration factor. Therefore, we expect the soluble ion concentrations to scale linearly at higher DMM concentrations until the solution becomes supersaturated or precipitation occurs, much like what happens when leaching physical simulants **(Appendix I)**. While there is some deviation from the target concentrations, all the macronutrient soluble species scale linearly with increased DMM concentrations up to 20x DMM **(Figure 1B)**. Therefore, we expect that using 20x DMM will still accurately simulate the soluble species present in the comparable leaching condition.

### Suggested Use Of DMM As A Microbial Growth Media

In **Box 1**, we describe recipes that translate the macro- and micro nutrient formulations (**Tables 2 and 3**) into salt mixtures that can be made from commercially available reagents.

Notably, these formulations do not include a carbon source. Methanogenic archaea^26^ or photosynthetic bacteria^89,90^ could utilize the CO_2_-rich atmosphere as a carbon source on Mars. Bicarbonates detected by SAM are naturally present in the Martian regolith and could also be added to DMM in this context. For heterotrophic bacteria, the CO_2_ atmosphere could be electrochemically fixed, or even biologically converted, into acetic acid, to serve as an aqueous ISRU-derived C_2_ carbon source^91–93^. Other electrochemically derived carbon sources, like ethanol, methanol, or formic acid, could also be used and may be preferable depending on a given microbe’s metabolism^94^. We chose acetate as the carbon source for our microbiology experiments to simulate heterotrophic growth in a habitable aqueous chemical environment on Mars.

Depending on the organism’s metabolism, growth might be impacted by the absence of a buffering agent, as metabolic activity can drive significant shifts in pH. DMM is formulated at a pH of 7.7, but pH often shifts during growth as nutrients are consumed. In M9, an often used chemically defined minimal medium for microbiology^95^, a high concentration of soluble phosphates acts as a buffering agent to maintain neutral pH during microbial growth. While phosphate buffering is common in biological studies, it would break the integrity of sourcing essential elements *in situ*.

In media lacking natural buffering, zwitterionic compounds with high solubility and pKa values around neutral pH, like Tris, HEPES, MOPS, or TES, are often added to maintain viable growth conditions^96,97^. In controlled bioreactor systems, pH is typically held constant by adjustment with acid or base addition. Bicarbonates in the regolith could serve as a potential ISRU-derived buffering agent. However, bicarbonate buffering is centered around a pKa of ∼ 6.3 and therefore provides limited buffering capacity near neutral pH. Because its effectiveness depends on equilibrium with CO_2_ and biological activity, it may not provide stable pH control under all experimental conditions. Alternatively, a buffering agent may not be needed if a microbe of interest is able to withstand pH changes during growth. In fact, it is often recommended to exclude buffering agents when conducting physiological characterization^98^. As a modifiable defined medium, **DMM** can incorporate any of these buffering options. We chose to utilize TES to maintain the target pH during growth experiments as needed, as it had the most minimal impact on measured growth compared to other buffering agents tested.

Based on our experimental characterization (**Figure 1)**, soluble ionic concentrations in DMM scale linearly from 1x to 20x DMM, allowing for simulating environments with higher input regolith to create richer media. Screening across a range of concentrations enables identification of conditions that provide sufficient nutrients while avoiding inhibition by soluble stressors. We can run growth assays in up to 20x DMM (i.e. 800 g of regolith / 1 liter of water). Creating even higher DMM concentrations can be informed by regolith dissolution experiments, based on future *in situ* measurements or available terrestrial simulants **(Appendix I**).

## APPLICATIONS

### Using DMM To Benchmark Microbes For Chemical Habitability

Next, we benchmarked standard lab and industry microbes for their Mars chemical habitability potential, as measured by growth in DMM. In DMM, strains have to utilize acetate (a Mars-relevant carbon source), rely on nitrate as the sole fixed nitrogen source, obtain essential trace elements, and tolerate any limiting amount of perchlorates, other salts, and/or metals. Using our defined media allows us to modify components, such as the sources of nitrogen, phosphorus, and carbon or the salt concentration to investigate the contributing factors to microbial survival and growth in DMM. Because strains able to survive and proliferate in DMM are more likely to grow in a Mars chemical environment, we can use DMM to investigate the potential for bio-ISRU.

We demonstrated investigating this by probing the key differences between M9, a common chemically defined minimal media, and DMM: carbon source, nitrogen source, phosphate concentration, presence of perchlorates, presence of trace elements, and buffering capacity **(Figure 2A)**. We consider M9 as the defined “Earth” laboratory conditions containing ammonium as a nitrogen source, high levels of bioavailable phosphate that also serve as a buffering agent, glucose or pyruvate as a preferred carbon source, the presence of solely essential trace elements, and lacking stressors like perchlorates. In contrast, DMM defines the “Mars” chemical environment containing nitrate as a nitrogen source, low levels of phosphate and no additional buffering, acetate as a carbon source, the presence of stressors like perchlorate and potentially toxic metals, in addition to essential trace elements sourced at low levels from the regolith.

**Figure 2:**
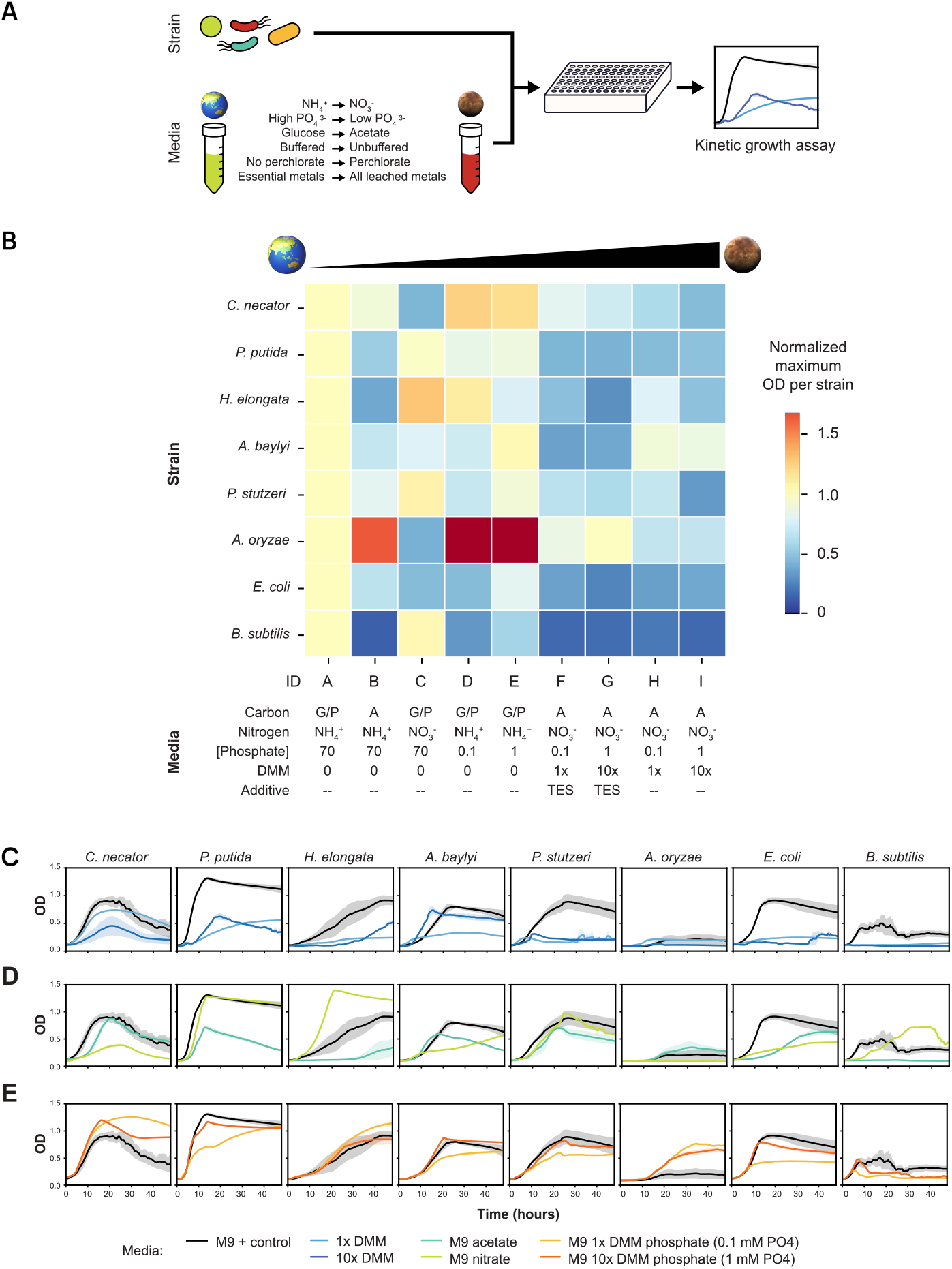
Benchmarking of strains-of-interest in DMM, and analysis of their limiting conditions. **A**. Experimental design for benchmarking microbial growth in a stepwise gradient of Earth-like to Mars-like conditions. We measured kinetic growth curves for each strain grown in various Earth-to-Mars media formulations in the 96-well plates. **B**. Heat map of the normalized maximum OD_600_ per strain in M9 minimal media with glucose or pyruvate as a carbon source. normalized (to OD_600_ = 1) and compared to increasingly Mars-like conditions. Media A-I are described fully in the Materials and Methods. **C**. Growth curves of strains grown in M9 (Media A. black) compared to lx (Media H. light blue) and 10x (Media I, dark blue) DMM. All curves have shaded regions indicating 95% confidence intervals. **D**. Growth curves of strains grown in M9 minimal media with glucose or pyruvate and ammonium (Media A, black line), with acetate and ammonium (Media 8, green line), or with glucose or pyruvate and nitrate (Media C, yellow line) as a carbon and a nitrogen source. All curves have shaded regions indicating 95% confidence intervals. **E**. Growth curves of strains grown in M9 minimal media with M9-levels of phosphate (Media A. black line) or lx (Media D. light orange) or 10x (Media E. dark orange) levels of phosphate. All curves have shaded regions indicating 95% confidence intervals. In panels B, C, D, and E, n = 2, 4, or 6; shown in more detail in **Supplementary Figure 1**.

We tested *Escherichia coli* MG1655, *Bacillus subtilis* PY79, *Pseudomonas putida* KT2440, *Pseudomonas stutzeri* AW-1, *Halomonas elongata* 1H9, *Cupriavidus necator* H16, *Azospira oryzae* PS, and *Acinetobacter baylyi* ADP1 for their ability to grow in aqueous growth media on a stepwise gradient from Earth-to Mars-like conditions (Figure **2B**). We aimed to screen a variety of bacteria, like standard lab strains *(E. coli*, B. subtilis) and relevant heterotrophic bacteria for Mars chemical habitability, such as halophiles *(H. elongata)*, perchlorate-reducing bacteria *(P. stutzeri, A*. oryzae), and biomanufacturing chassis *(P. putida*, C. *necator, A. baylyí*). In moving from Earth-like to Mars-like conditions, we tested individual reagent substitutions to investigate which factors may limit growth in full DMM conditions (ex. comparing the growth of strains in M9 with ammonium (Figure 2B, Media A) as a fixed nitrogen source to M9 with nitrate (Figure 2B, Media C)) under aerobic, 30°C conditions. Many strains showed growth in the most Mars like chemical environment, though the maximum optical density (max OD), growth rate, and lag times depended on the exact media conditions. The top-performing strains (C. *necator, P. putida, H. elongata*, and *A. baylyí*) were able to readily utilize nitrate and acetate, and withstand increasing concentrations of DMM, while maintaining reasonable levels of growth.

Compared to growing in M9 with a preferred carbon source (pyruvate for C. *necator* and glucose for all other strains), all strains experience a reduction of 34% to 90% in their max OD when grown in lx or 10x DMM with acetate as a carbon source (Figure 2C). In some instances, like for *A. baylyi*, growth in lx DMM is lower than in 10x DMM suggesting possible nutrient limitations at lower regolith to water ratios. To investigate this further, we explored the individual effects of carbon and nitrogen sources, and phosphate concentrations.

As acetate is a Mars-relevant carbon source, we compared growth in M9 with glucose or pyruvate to M9 with acetate (Figure 2D). While all strains except B. *subtilis* were able to utilize acetate, most experienced significant decreases in their max OD. Only C. *necator* performed better with acetate than with its positive control carbon source, pyruvate. For the differences in fixed nitrogen source, some strains had a lower max OD utilizing nitrate *(E. coli, A*. oryzae), while other strains grew to the same (P. *putida, P. stutzen)* or higher (B. *subtilis, H. elongata)* max OD with nitrate rather than ammonium (Figure 2D). While low phosphate levels have a diminishing effect on max OD in M9 minimal media with “lx DMM amount” of phosphate (0.1 mM). there is sufficient phosphate supplied in “10x DMM amount” (1 mM) relative to growth in M9 minimal media with standard phosphate levels and buffering (Figure 2E).

### Sensitivity Analysis To Micronutrients And Toxins In DMM

To evaluate the utility of the DMM micronutrient portion, we conducted sensitivity analysis to determine how differences between DMM and actual Martian chemistry, stemming from site-to-site variation or measurement uncertainty, meaningfully impact microbial growth. Mars is broadly basaltic, but local mineralogical heterogeneity exists across the planet with enrichment or depletion of certain chemical species^74,80,99,100^. The composition of the top layer of fine dust is thought to be quite homogenous due to frequent global dust storms. while larger particle sized granules exhibit more heterogeneity^80,101^. DMM’s micronutrient formulation similarly carries uncertainty, given its basis on terrestrial analog leachate data due to the absence of *in situ* aqueous data on trace elements, and because trace elements can be growth-limiting at both deficiency and toxicity thresholds depending on the microbe and growth environment. The flexibility of a defined media allows us to investigate the potential impact of uncertainty in the measured concentrations of soluble micronutrients by studying growth robustness in modified DMM.

We first compared DMM’s trace metal concentrations to published essentiality and toxicity thresholds for each element. Using available data from the literature on metal toxicity in *E. coli*, an extensively studied model bacterium, we see that no soluble micronutrients in 1x DMM reach a lethal concentration and most are several orders of magnitude lower (Figure 3A). Understanding an organism’s nutrient requirement is more difficult than toxicity. Research quantifying the intracellular concentrations of trace metals or growing cells in metal depleted cultures serves as a useful reference point for understanding if order-of-magnitude micronutrient concentrations in DMM may limit growth. Available data on *E. coli* highlights that the measured soluble concentrations of some trace metals in DMM may be limiting for growth. For example, while 1x DMM provides adequate iron, the quantities of copper, zinc, nickel, manganese, or molybdenum may be insufficient for growth compared to *E*. co/i’s chemical habitability zone, calculated using the reported nutrient requirement and inhibitory concentrations from literature (Figure 3A).

**Figure 3:**
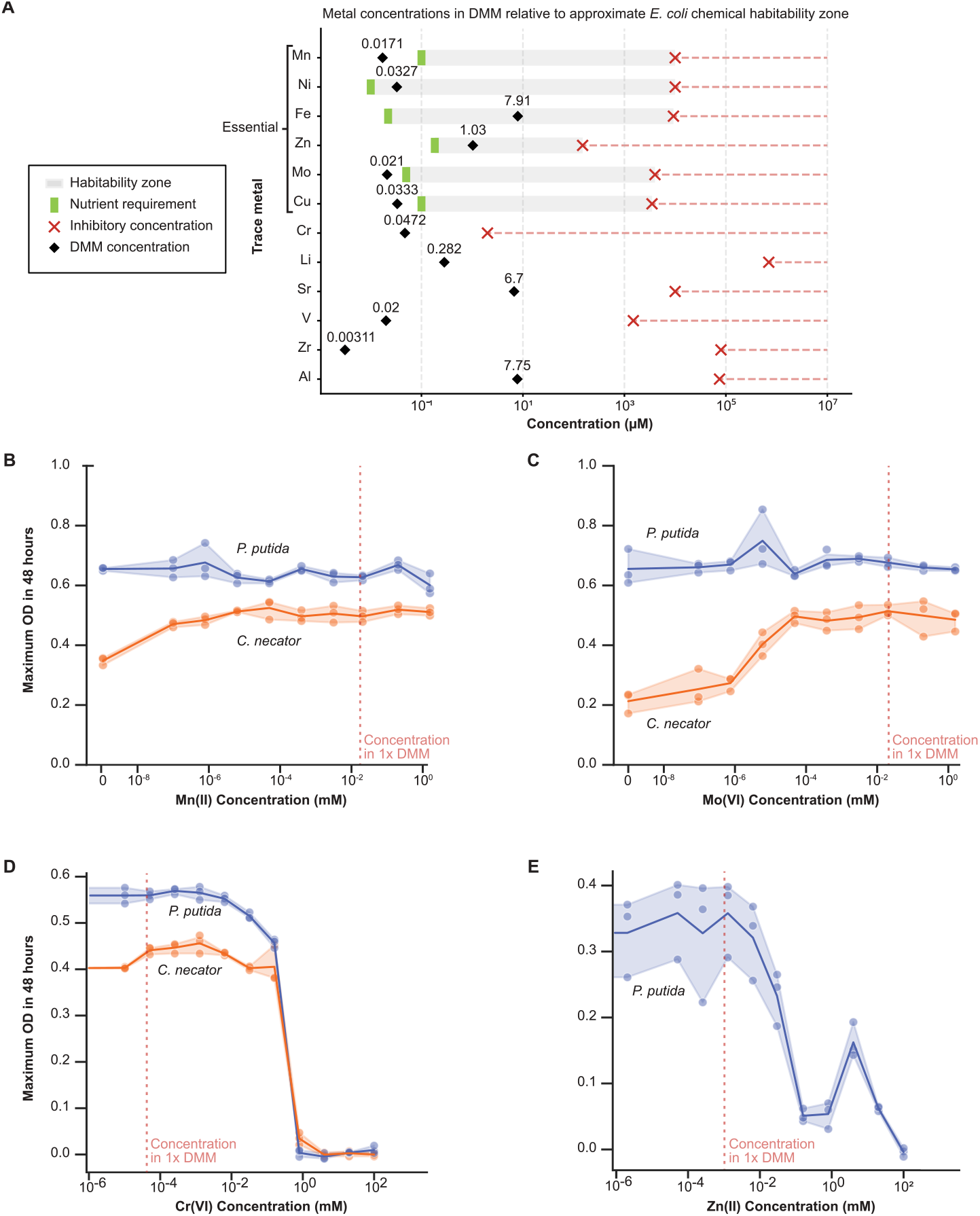
Sensitivity analysis to potential DMM deviations. **A**. Comparison of soluble concentration of trace elements in lx DMM to nutrient requirement and inhibitory concentrations of those trace elements for *E*.*coli* reported in the literature (Supplementary Table 3) **B**. Maximum OD_600_ values reached in 48 hours for *P. putida* (blue line) and C. *necator* (orange line) grown in 10x DMM with 0-2 mM Mn(II). The reference line indicates concentration of Mn(II) in DMM micronutrient formulation. Shaded region indicates range between replicates (n = 3). **C**. Maximum OD_600_ values reached in 48 hours for *P putida* (blue line) and C. *necator* (orange line) grown in 10x DMM with 0-2 mM Mo(VI). The reference line indicates concentration of Mo(VI) in DMM micronutrient formulation. Shaded region indicates range between replicates (n = 3). **D**. Maximum OD_600_ values reached in 48 hours for *P putida* (blue line) and C. *necator* (orange line) grown in 10x DMM with 0-100 mM Cr(VI). The reference line indicates concentration of Cr(III) in DMM micronutrient formulation. Shaded region indicates range between replicates (n = 3). **E**. Maximum OD60o values reached in 48 hours for *P putida* grown in 10x DMM with 0-700 mM Zn(II). The reference line indicates concentration of Zn(II) in DMM micronutrient formulation. Shaded region indicates range between replicates (n = 3).

We performed sensitivity analysis to micronutrient concentration by measuring microbial growth in dose-response assays with DMM and specific micronutrients. To demonstrate determining the potential growth limiting impact of DMM’s micronutrient formulation, we chose to look at the nutrient requirement of molybdenum and manganese for two promising strains from the benchmarking experiments (P. *putida* and C. *necator)*. Molybdenum (Mo) is of interest because it acts as a metal cofactor for nitrogen fixation and reduction for almost all diazotrophs^69,102^, processes that are significant given that nitrate is the exclusive source of fixed nitrogen in DMM. Manganese (Mn) plays an important role in oxidative stress resistance, which is of interest as DMM contains perchlorate, which causes oxidative stress^103-105^. Also of interest, accumulation of manganese has been seen as a key defense mechanism for radiation and oxidative stress damage in Oeinococcus radiodurans^106,107^ and higher desiccation tolerance in dry-climate soil bacteria^108^.

When testing growth in Mn or Mo limiting conditions, P. *putida* continued to have robust growth in 10x DMM, even without it supplied (Figure 3B, 3C). *C. necator* was able to grow in 10x DMM with no supplied Mn, but showed improved growth when a minimal amount of Mn (∼1x 10^-7^ mM) was supplied (Figure 3B). In the more limiting Mo conditions (0 - 6 x 10^-6^ mM), C. *necator* had poorer growth in 10x DMM, characterized by lower growth rates and lower maximum 0D600 reached in 48 hours (Figure 3C). C. *necator* was able to grow uninhibited in 10x DMM when Mo was supplied above ∼6 x 10^-6^ mM. When determining the potential of DMM supplying inadequate trace nutrients for growth, we see that the soluble Mn and Mo concentrations in DMM are approximately one and four orders of magnitude higher respectively than the limiting concentrations determined in this experiment for these two microbes. Other trace elements at low concentrations, such as copper and nickel, may still be limiting due to uncertainties in Martian regolith data and should be investigated independently for other organisms of interest.

To demonstrate the potential impact caused by high concentrations of cytotoxic species in DMM’s micronutrient formulation, we investigated the toxicity effect of chromium and zinc. Chromium is an often cited potential hazard on Mars, as hexavalent chromium (Cr(VI)) is a known human carcinogen^84,109,110^. Depending on the microorganism, chromium can be a potential stressor, yet there are many naturally occurring bacteria that can reduce Cr(VI) to the less harmful Cr(III), such as P. putida^88,111-113^ While zinc (Zn) is an essential trace metal, zinc at excessive concentrations can inhibit microbial growth by interacting with sulfhydryl side groups of proteins and repressing dehydrogenase activity^114-116^.

We performed dose-response growth assays for C. *necator* and P. *putida* in 10x DMM containing up to 100 mM Cr(VI) or Zn. Both strains saw inhibited growth between 0.16-0.8 mM Cr(VI), as seen by little to no growth after 48 hours measured by optical density (Figure 3D). When P. *putida* was grown in 10x DMM containing a range of Zn concentrations, growth started to be inhibited in media with 0.03 mM Zn and growth was completely inhibited at 0.2 mM Zn (Figure 3E). We chose not to run a similar test for C. *necator* as C. *necator* is expected to have three times higher tolerance for zinc than P. putida^117,118^. For the potential cytotoxic nature of DMM micronutrients, we see that the soluble Zn concentration in DMM is at least two orders of magnitude lower than the inhibitory concentration measured here for P. *putida* (Figure 3E). While DMM does not contain Cr(VI), the concentration of Cr(III) in DMM is approximately four orders of magnitude lower than the inhibitory concentration of Cr(VI) (Figure 3D). Together, these sensitivity analyses can demonstrate that DMM’s micronutrient concentrations pose no meaningful toxicity risk to the tested organisms while remaining at or above growth-permissive thresholds, establishing a useful baseline for systematically evaluating microbial habitability in Mars-relevant chemical conditions.

## DISCUSSION

Here we report our Defined Mars Media (DMM), a chemically defined Mars regolith extract analog designed for astrobiology experiments. We formulated DMM with available *in situ* data, using Curiosity rover and Phoenix lander data, along with literature equilibrium models, to formulate the major soluble species. Without *in* situ aqueous data on trace elements, we turned to using in-house leachate data of commercial physical Mars regolith analogs to formulate an optional micronutrient mix. As opposed to using traditional microbiology trace element mixes (ex. Wolfe’s mixture^119^ or SL-10^120^), DMM micronutrients allows us to incorporate elements rooted in both biological relevance and Martian geochemistry. Functionally, we recommend its use as a 40-800 g/L regolith:water leachate simulant that contains chemically defined quantities of soluble minerals. The defined formulation allows users to tune the concentrations of chemical species-of-interest to probe chemical habitability limitations such as perchlorate tolerance or to explore compatibility with different metabolisms. Depending on the intended use case, we suggest ways of adapting it to different Mars-relevant carbon sources and buffering conditions. Supplemental chemicals, like EDTA and TES, are added to improve experiment setup without impacting chemical habitability results. In contrast to existing Mars regolith analogs, DMM enables high fidelity, modular, and reproducible aqueous experimentation agnostic of simulant leaching conditions (Appendix I). It is the first chemically defined Mars media that contains all major soluble species in regolith, including nutrients and stressors.

The DMM recipe consistently recapitulates our desired formulation across different users and experiments, highlighting its capability to translate between labs and research. Optimized ion chromatography methods will enable more rigorous QC monitoring of perchlorate and phosphate concentrations. By modifying the defined media, we demonstrate its use for benchmarking microbial chemical habitability in leached Mars regolith and identify Mars-relevant carbon source utilization, nitrate utilization, tolerance at higher DMM concentrations, and buffering capacity as common limiting factors for microbial growth. This work can provide a framework for prioritizing research directions for microbes for Mars, such as determining their preference for available nutrients (nitrate, acetate) and tolerance to stressors (perchlorate, limited phosphate). We show that C. *necator*, P. *putida, H. elongata, and A. baylyi* are able to grow using only ISRU-derived nutrients. Interestingly, we find that several of the tested microbes are able to tolerate the elevated levels of perchlorate in 10x DMM (∼30 mM), indicating perchlorates’ baseline presence in regolith is only one of the stressors in chemical habitability on Mars.

We further analyze our recipe with respect to specific potential limiting and toxic trace metals to ensure growth performance if DMM concentrations are not representative of actual regolith. We find that two promising strains, P. *putida* (an aerobic chemoorganoheterotroph) and C. *necator* (a facultative chemolithoautotroph), are not sensitive to manganese limitation when grown in DMM. Limiting amounts of molybdenum are potentially concerning for C. *necator*, in that this metal is within an order of magnitude of the limiting concentration we measured. We also find that P. *putida* experiences zinc toxicity and hexavalent chromium sensitivity, but at quantities that are orders of magnitude beyond what microbes would likely encounter in Martian regolith. For bio-ISRU studies, the current micronutrient formulation in DMM is sufficient to enable microbial growth without concerns of toxicity or nutrient limitation, and can be easily modified to investigate any microbe’s trace element requirements. The micronutrient formulation, while based on leached simulants where the trace mineral content is not a part of the intrinsic simulant design, can still serve as an initial baseline that incorporates additional Mars-relevant elements beyond standard trace elements. This kind of sensitivity analysis was not possible with prior leachate-based media, and is thus a major advantage of using a defined media.

DMM fills a critical gap for a high fidelity and reproducible media for use in microbiological studies, and particularly for aqueous high throughput experimentation. However, the commercially available Mars regolith analogs and leachates may still provide a better basis for hydrated soils, particulate radiation shielding, and plant root propagation experiments, as the solid phase plays a more important role in plant behavior90. Studies on biology’s interaction with carbonates may also benefit from simulating with solid media or a representative CO_2_ atmosphere, which is outside the scope of a soluble defined media. Evaluating chemical habitability in physical environments more similar to Mars (i.e. temperature, pCO_2_) can further inform future DMM formulations and allow for soluble bicarbonate in the simulated medium. Meanwhile, having a stable defined recipe makes it simple to prepare variants that take into account altered leaching conditions based on simulant experiments (ex. acidic leaching) or to customize the formulation to locations on Mars with known mineral enrichment. Future *in situ* measurements or analysis of returned regolith samples would allow researchers to fully characterize the composition and variability of a slew of different, but proximal, regolith sample sites. Prospective rover and lander missions could prioritize WCL-like experiments with expanded quantitative equipment so as to specifically measure for the bioavailable essential nutrients like nitrates, phosphates, sulfates, and trace micronutrients. These studies would also obtain first-of-its-kind information on the solubility and charge state of trace minerals, both biologically essential trace elements and putative heavy metal stressors. On Earth, solubility modeling of different mineral compositions on Mars would identify the upper and lower bounds for accessible nutrients, salts, and toxins when the ratio of regolith and water is adjusted to meet experimental needs. For example, DMM formulation may need to take into account species extractability when leaching at higher concentrations of regolith (Appendix II).

To increase accessibility and encourage larger adoption, we have onboarded a version of DMM at a 20x concentration (e.g. 800 g/L) at Space Resource Technologies (spaceresourcetech.com/collections/martian-simulants/products/dmm-defined-mars-media), the supplier for several of the physical regolith simulants used in this study. This purchasable media contains the macro- and micronutrient species, except perchlorate salts which need to be supplied by the user. Researchers interested in customizing DMM to fit their experiment or microorganism needs are encouraged to follow our instructions in **Box 1**, and can also reference the catalog numbers for the reagent ingredients in **Supplementary Table 1**. We made an effort to ensure each chemical is readily available from typical laboratory reagent distributors. We believe this chemically defined Mars leachate simulant will enable lab research in astrobiology (chemical habitability studies exploring the possibility of extant and/or extinct Martian life), biological ISRU on robotic missions, bioremediation, and terraforming research.

## MATERIALS AND METHODS

### Preparing Leachates Of Mars Regolith Simulants For IC And ICP-MS

The regolith simulants were weighed out in sterile 50 mL plastic conical tubes depending on the final desired concentration of solid materials. The leaching solutions were added to the solid materials in the conical tubes to a final volume of 50 mL. Deionized (DI) water was used for a neutral leachant and the non-neutral leaching solutions were prepared by diluting 1 M NaOH down to 100 mM NaOH (pH 12) and to 0.03 mM NaOH (pH 9) or by diluting 1 M HCI down to 100 mM HCI (pH l) and to 0.02-7 mM HCI (pH 5).

To start the leaching process. the tubes were shaken to fully resuspend the solid phase. The tops of the tubes were wrapped with Parafilm to prevent leaking. The tubes were set on their side and fastened to a microplate shaker (Fisherbrand Incubating Microplate Shaker, Catalog No. 02-217-757). The shaker was set to 350 rpm and 3 mm orbital throw. Tubes were shaken for 24 hours for standard leaching and up to 5 days when studying the impact of length of leaching. After the desired leaching time, the leachate was separated by centrifuging the tubes for 2 minutes at 2,500 x g at room temperature and decanting the supernatant into a sterile 15 mL tube. The leachates were sent to an external laboratory for ion chromatography (IC), inductively coupled plasma optical emission spectroscopy (ICP-OES) and inductively coupled plasma mass spectrometry (ICP-MS).

### Preparing Different Stock Concentrations Of Defined Mars Media (DMM)

A 20x concentrated stock of DMM was prepared according to the protocol and recipe described in **Box 1**. Briefly, three stock solutions were prepared, representing a 500x bulk salts solution (23.08 g/L NaCl, 25.28 g/L KNO_3,_21.25 g/L NaNO_3_, 164.48 g/L Na_2_SO_4_· 10H_2_O, 367.13 g/L MgSO_4_ · 7H_2_O). 1000x perchlorate solution (91.11 g/L Mg(CIO_4_)_2_ · 6H_2_O, 362.92 g/L Ca(CIO_4)2_ · 4H_2_O) and 1000x phosphate solution (18.23 g/L Na_2_HPO_4_ · 7H_2_O, 4.42 g/L NaH_2_PO_4_ · H_2_O). The stock solutions were diluted in deionized water to a 20x final concentration, representing 20x DMM. Aliquots of the 20x DMM solution were further diluted down to 10x, 8x, 5x, and lx concentrations. In addition, a lx DMM solution was prepared similarly, by diluting down stock solutions to a final lx concentration. After sterile filtration, all solutions were sent for elemental and ionic analysis. To understand recipe consistency, the same solutions were prepared by a second operator and also sent for elemental and ionic analysis.

### Elemental And Ionic Analysis (IC, ICP-OES, ICP-MS)

All IC, ICP-MS, and ICP-OES measurements were done at the Center for Applied Isotope Studies at the University of Georgia (www.cais.uga.edu). in the Laboratory for Environmental Analysis (LEA) for IC and Plasma Chemistry Laboratory for ICP-MS and ICP-OES.

Anion chromatography was used to determine the soluble concentrations of Cl^-^(chloride), NO_3_ ^-^ (nitrate), PO_4_^3-^ (phosphate), SO_4_^2-^ (sulfate), and CIO_4_^-^ (perchlorate). Ion chromatography was performed using a Dionex GP40 IC system with a Thermo Scientific Dionex ADRS 600 and Dionex ED40 as the suppressor and electrochemical detector respectively. The typical method used was pumping samples over a Dionex lonPac AS20 column with an AG22 guard column attached. The mobile phase was 4.5 mM sodium carbonate and 1.4 mM sodium bicarbonate. The run was conducted for 40 minutes at 0.8 mL/min flow rate. Elution times of 7.0, 11.1, 15.2, 17.6, and 34.l were used for chloride, nitrate, phosphate, sulfate, and perchlorate respectively. For some older samples submitted, a method using an lonPac AS16 column with a mobile phase of 12.5-18.5 mM NaOH was used but resulted in more interference when measuring phosphate concentrations.

Elements chosen to be measured via ICP-MS or ICP-OES belong to three categories: biologically relevant elements (Na, K, Mg, Ca, Mn, Cu, W, B, Fe, Zn. Ni. Mo, Co, Se), elements included in oxides that make up simulants (ex. Si, Ti. Al), and other Mars-relevant elements previously detected at high concentrations compared to Earth (ex. Cr, Zr, Sr). We determined which biologically relevant trace elements to include in elemental analysis by looking at commonly used trace element mixes, like Wolfe’s mixture^119^ or SL-10^120^· which are available through typical reagent suppliers like ATCC, ThermoFisher. or Sigma. The chemical make-up of the commercial regolith simulants is available on the supplier websites^121,122^.

Elemental analysis was performed using ICP-MS and ICP-OES. Samples and method blanks were diluted in metal-free polypropylene centrifuge tubes using trace metal grade acids: 2% HNO_3_ for measurement of Na, K, Ca, Mg. B, Li, Be, Al, Si, Ti, V, Cr, Mn, Fe, Co, Ni, Cu, Zn, As. Se, Sr, Zr, Ag. Te, Cs, Ba, W, Pb, Th, and U; or 5% w/w HCI for Mo, Sn, Sb, and Ta. The concentrations of Li, Be, Mg. Al, K, Ti, V, Cr, Mn, Fe, Co, Ni, Cu, Zn, As. Se, Sr, Zr, Ag. Mo, Sn, Sb, Te, Cs, Ba, Ta, W, Pb, Th, and U were determined by ICP-MS. A Thermo X-Series 2 ICP-MS with collision cell technology (CCT) and chilled spray chamber was used in kinetic energy discrimination (KED) mode with a mixture of 8 % hydrogen in helium to reduce polyatomic interferences for all elements except Li, Be, and U. The concentrations of B, Ca, Na, and Si were determined by ICP-OES. A Perkin Elmer Optima 8300 Dual View was used in radial mode for all elements to reduce interferences except B and Si, for which a deconvolution technique was used to subtract interferences. For both ICP-OES and ICP-MS, In was supplied inline as the internal standard for drift correction. All measured concentrations and limits of quantitation (LOQs) were corrected for dilution factors and any results below the LOQ for each element were reported as less than the LOQ.

### Preparing Microbial Growth Media For Benchmarking Assays

We prepared M9 minimal media as described by Cold Spring Harbor Protocols^95^· We made stocks of 5x M9 salts, 5x M9 salts without phosphate salts, 5x M9 salts without ammonium salts, 5x potassium nitrate at equimolar nitrogen to that found in 5x M9 salts, 1000x DMM phosphates. 5x 20% w/v glucose. 20% w/v sodium pyruvate, 20% w/v potassium acetate, lM MgSO_4_, lM CaCl_2_, and 20x DMM. From these. we were able to prepare the nine media in which strains were benchmarked as shown in **Figure 2A**. These media are as follows: A) M9 with 1% glucose. B) M9 with 1% acetate, C) M9 with 1% glucose and nitrate instead of ammonium, D) M9 with 1% glucose and lx DMM phosphates (0.1 mM phosphate). E) M9 with 1% glucose and 10x DMM phosphates (l mM phosphate). F) lx DMM with 1% acetate and 50 mM TES, G) 10x DMM with 1% acetate and 50 mM TES, H) lx DMM with 1% acetate, and I) 10x DMM with 1% acetate. For C. *necator*, in all instances where 1% glucose was used, 1% pyruvate was also added.

### Growth Assays Of Bacteria In DMM

We obtained the following bacterial strains from the American Type Culture Collection (ATCC). Bacillus Genetic Stock Center (BGSC), Coli Genetic Stock Center (CGSC), and Leibniz Institute DSMZ-German Collection of Microorganisms and Cell Cultures (DSMZ): *Acinetobacter baylyi* ADP1 (DSM 24193). *Azospira oryzae* PS (DSM 13638). *Bacillus subtilis* PY79 (BGSCID 1A747), *Cupriavidus necator* H16 (DSM 428), *Escherichia coli* MG1655 (CGSC #6300), *Halomonas elongata* 1H9 (ATCC 33773), *Pseudomonas stutzeri* AW-1 (DSM 13592), and *Pseudomonas putida* KT2440 (DSM 6125). We grew them up in their recommended media and glycerol stocked them for long-term -80°C storage.

For benchmarking the strains in M9 and DMM media, we first started overnights of the strains in rich media: Luria Broth (LB) (most strains), Super Optimal Broth (SOB) (C. *necator)*, LB + 3% NaCl (H. *elongata)*, or ALP minus lactate *(A*. oryzae). After 76-20 hours of growth at 30’C, the OD_600_ values of strains were measured. The benchmarking media set was prepared as described previously, and 700 µL was aliquoted out into a Thermo Scientific Nunc 96-Well Optical Bottom Microplate (Fisher Scientific 12-566-70). 5 µL of overnight culture normalized between 0.3 and 0.7 OD_600_ nm was added to the wells and growth curves were conducted for 72 hours at 30°C with high speed shaking and readings every 70 minutes, in either a BioTek Synergy HT or a BioTek Synergy Mx plate reader.

For dose-response growth assays studying trace element concentration sensitivity, 10-15x DMM + 1-1.5% potassium acetate (Fisher Scientific, AAA1632136) + 50 mM TES pH 8 (Fisher Scientific, AAB2181918) was prepared as described above. For each trace element being tested, a serial dilution was prepared to test DMM media with 0-100 mM of the trace metal included. If a 0 mM metal condition was tested, the micronutrient mix was remade with that specific metal removed. Cultures of *P. putida* and C. *necator* were prepared by growing overnight in SOB media (28 g/L SOB powder in DI water; Fisher Scientific DF0443-17) to a final OD_600_ of 6-8. Cultures were pelleted and resuspended in a solution of 1% NaCl + 10 mM TES pH 8. Pellets were washed three times in the same solution and normalized to an OD_600_ of 1. A 96-well black assay plate (Fisher Scientific, 72-566-70) filled with 100 µL of media was inoculated with 3 µL of normalized culture. Growth curves were conducted for 48-72 hours at 30°C with high-speed shaking and readings every 10 minutes, in either a BioTek Synergy HT or a BioTek Synergy Mx plate reader.

## ACKNOWLEDGEMENTS

This research was funded by The Astera Institute. We would like to thank Michael **H**. Hecht for extensive guidance and serving as a reviewer. We would also like to thank Maria-Paz Zorano, Anna Metke, Edwin Kite, Alfonso Davila, Mohit Melwani Daswani, Suniti Karunatillake, and Susanne Schwenzer for advice and feedback throughout this research effort. Rachel Harris is supported by an appointment to the NASA Postdoctoral Management Program through the Astrobiology Program at NASA Headquarters, administered by Oak Ridge Associated Universities under contract with NASA. We thank Alfonso Davila and Samuel Kounaves for helping us obtain original batches of MGS-1, JEZ-1, and MMS-1 regolith simulants. We thank scientists from the University of Georgia’s Center for Applied Isotope Studies: Dr. Sarah C. Jantzi (Plasma Chemistry Laboratory); Dr. Sayed Hassan and Dr. Tret Burdette (Laboratory for Environmental Analysis) for the elemental and ionic analyses and guidance. We thank Rachel Sevey and Olesia Bushkova for their assistance with scientific communication. We thank Anna Metke at Space Resource Technologies for working with us to set up DMM as a commercially available Mars regolith simulant. Lastly, we thank our external peer reviewers from Research Hub for providing feedback that was incorporated into the subsequent version of the manuscript. Peer reviews can be viewed at www.researchhub.com/paper/11204937.

## SUPPLEMENTARY INFORMATION

**Supplementary Table 1:** Reagents Used In Preparing DMM In-House.

| Supplementary Table 1: Reagents Used In Preparing DMM In-House. |  |  |  |  |
| --- | --- | --- | --- | --- |
| Chemical Name | Formula | Manufacturer | Catalog Number | CAS |
| Sodium chloride | NaCl | Thermo Fisher | BP358 | 7647-14-5 |
| Potassium nitrate | KNO<sub>3</sub> | Thermo Fisher | P263 | 7757-79-1 |
| Sodium sulfate decahydrate | Na<sub>2</sub>SO<sub>4</sub> · 10H<sub>2</sub>O | Thermo Fisher | AAA157020B | 7727-73-3 |
| Magnesium sulfate heptahydrate | MgSO<sub>4</sub> · 7H<sub>2</sub>O | Thermo Fisher | AA1159636 | 10034-99-8 |
| Sodium nitrate | NaNO<sub>3</sub> | Thermo Fisher | S343 | 7631-99-4 |
| Calcium perchlorate tetrahydrate | Ca(ClO<sub>4</sub>)<sub>2</sub> · 4H<sub>2</sub>O | Thermo Fisher | AC390631000 | 15627-86-8 |
| Magnesium perchlorate hexahydrate | Mg(ClO<sub>4</sub>)<sub>2</sub> · 6H<sub>2</sub>O | Sigma Aldrich | 309303 | 13446-19-0 |
| Disodium hydrogen phosphate heptahydrate | Na<sub>2</sub>HPO<sub>4</sub> · 7H<sub>2</sub>O | Thermo Fisher | S373-500 | 7782-85-6 |
| Sodium dihydrogen phosphate monohydrate | NaH<sub>2</sub>PO<sub>4</sub> · H<sub>2</sub>O | Sigma Aldrich | S9368 | 10049-21-5 |
| Aluminium potassium sulfate dodecahydrate (Potassium alum) | AlK(SO<sub>4</sub>)<sub>2</sub> · 12H<sub>2</sub>O | Thermo Fisher | AC423260050 | 7784-24-9 |
| Iron(II) sulfate heptahydrate (Ferrous sulfate) | FeSO<sub>4</sub> · 7H<sub>2</sub>O | Thermo Fisher | AC423731000 | 7782-63-0 |
| Strontium chloride hexahydrate | SrCl<sub>2</sub> · 6H<sub>2</sub>O | Thermo Fisher | AC315081000 | 10025-70-4 |
| Zinc sulfate heptahydrate | ZnSO<sub>4</sub> · 7H<sub>2</sub>O | Sigma Aldrich | Z0251 | 7446-20-0 |
| Barium chloride dihydrate | BaCl<sub>2</sub> · 2H<sub>2</sub>O | Thermo Fisher | AA1064614 | 10326-27-9 |
| Boric acid | H<sub>3</sub>BO<sub>3</sub> | Sigma Aldrich | B6768 | 10043-35-3 |
| Sodium molybdate dihydrate | Na<sub>2</sub>MoO<sub>4</sub> · 2H<sub>2</sub>O | Sigma Aldrich | 331058 | 10102-40-6 |
| Lithium chloride | LiCl | Thermo Fisher | AC413270050 | 7447-41-8 |
| Chromium(III) chloride hexahydrate | CrCl<sub>3</sub> · 6H<sub>2</sub>O | Thermo Fisher | AA1068606 | 10060-12-5 |
| Nickel(II) chloride hexahydrate | NiCl<sub>2</sub> · 6H<sub>2</sub>O | Sigma Aldrich | 654507 | 7791-20-0 |
| Copper(II) chloride dihydrate | CuCl<sub>2</sub> · 2H<sub>2</sub>O | Sigma Aldrich | 459097 | 10125-13-0 |
| Manganese(II) chloride tetrahydrate | MnCl<sub>2</sub> · 4H<sub>2</sub>O | MP Biomedicals | ICN15533480 | 13446-34-9 |
| Vanadyl sulfate hydrate | VO<sub>2</sub>SO<sub>4</sub> · nH<sub>2</sub>O | Sigma Aldrich | 204862 | 123334-20-3 |
| Sodium selenite | Na<sub>2</sub>SeO<sub>3</sub> | Sigma Aldrich | 214485 | 10102-18-8 |
| Sodium tungstate dihydrate | Na<sub>2</sub>WO<sub>4</sub> · 2H<sub>2</sub>O | Thermo Fisher | AC207620050 | 10213-10-2 |
| Cobalt(II) chloride hexahydrate | CoCl<sub>2</sub> · 6H<sub>2</sub>O | Sigma Aldrich | C8661 | 7791-13-1 |
| Zirconium(IV) sulfate tetrahydrate | Zr(SO<sub>4</sub>)<sub>2</sub> · 4H<sub>2</sub>O | Thermo Fisher | AA4319414 | 7446-31-3 |
| Silver nitrate | AgNO<sub>3</sub> | Thermo Fisher | AC197680050 | 7761-88-8 |

**Supplementary Table 2.** Martian regolith simulants used in generating DMM micronutrient mix (Table 3) and in leaching study (Appendix I). Current batches were purchased from vendors in April 2025. Older batches were kindly gifted by Alfonso Davila, for MGS-1 and JEZ-1, and Samuel Kounaves, for MMS-1.

| Simulant | Vendor | Description | Batch Number |
| --- | --- | --- | --- |
| Enhanced Mars Simulant (MMS-2)<sup>33, 122</sup> | The Martian Garden (Austin, TX) | Follow-up to Mojave Mars Simulant, originally made to support Mars Phoenix lander development | Current. MMS-2 Purchased April 2025<br>Old. MMS-1, from JPL, 2008 |
| Mars Global Simulant (MGS-1)<sup>37</sup> | Space Resource Technologies (Oviedo, FL) | Several terrestrial minerals mixed to be mineralogical standard analog based on Curiosity rover data | Current. 003-05-001-0523<br>Old. 001-05-001-0120 |
| Jezero Delta Simulant (JEZ-1)<sup>37</sup> | Space Resource Technologies (Oviedo, FL) | Jezero Crater delta analog based on Perseverance rover data | Current. 002-08-001-0621<br>Old. 001-08-001-0120 |

**Supplementary Figure 1.**
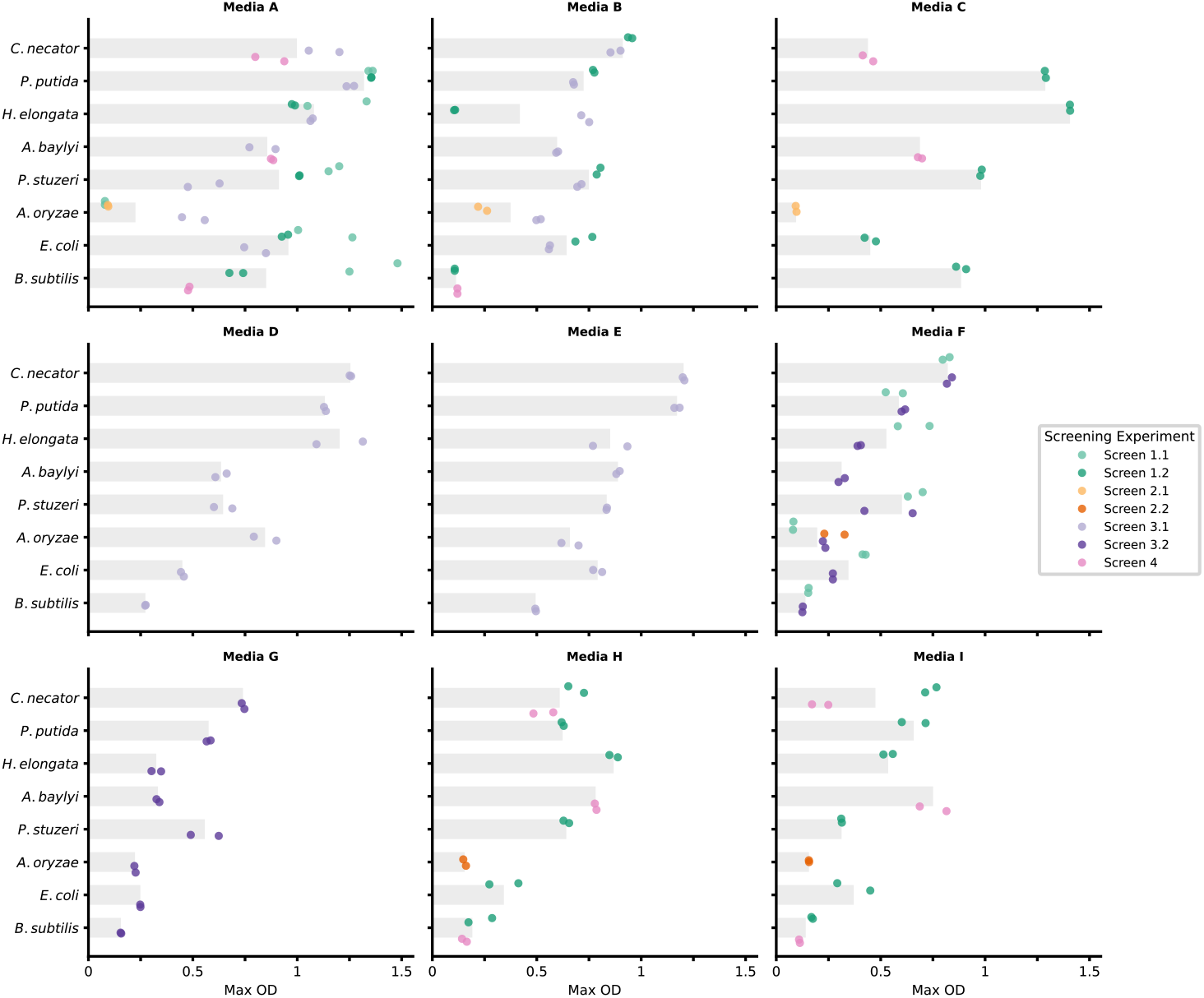
Non-normalized maximum OD per strain in the Earth-to-Mars media formulations. Barplots of the maximum ODs from kinetic growth curves of strains grown in different Earth-to-Mars media formulations with duplicate measurements per experiment colored by the benchmarking screening experiment they are a part of. Every screened strain and media combination has n = 2 to n = 6 replicates depending on how many benchmarking screening experiments the pair is a part of. Media A-I are described fully in the Materials and Methods.

**Supplementary Table 3.** Nutrient requirement and inhibitory concentration values of trace metals in studies of *E. coli*, a Gram-negative heterotrophic bacteria.

| Trace Element | Values Used In Figure 3A And Supplementary Figure 5B | References |
| --- | --- | --- |
| Manganese | **Nutrient Requirement:** 0.01 µM Intracellular Mn (measured via ICP-MS) | 103, 123 |
|  | **Inhibitory Concentration:** 10 mM Mn(II) | 124 |
| Nickel | **Nutrient Requirement:** 0.07 µM Ni (measured via ICP-MS) | 123, 125 |
|  | **Inhibitory Concentration:** 10 µM Ni(II) | 126 |
| Iron | **Nutrient Requirement:** 0.022 µM Fe | 127, 128 |
|  | **Inhibitory Concentration:** 9.25 mM Fe(III) / 11.25 mM Fe(II) | 129 |
| Zinc | **Nutrient Requirement:** 0.183 µM Zn (measured via ICP-MS) | 130 |
|  | **Inhibitory Concentration:** 0.15 mM Zn(II) (complete inhibition at 0.35mM) | 115 |
| Molybdenum | **Nutrient Requirement:** 0.05 µM Intracellular Mo (measured via ICP-MS) | 131 |
|  | **Inhibitory Concentration:** 4 mM Mo(VI) | 132 |
| Copper | **Nutrient Requirement:** 0.1 µM Cu(II) | 133, 134 |
|  | **Inhibitory Concentration:** 3.5 mM Cu(II) | 135 |
| Chromium | **Inhibitory Concentration:** 20 µM Cr(III) / 100 µM Cr(VI) | 136, 137 |
| Lithium | **Inhibitory Concentration:** 0.7 M Li(I) | 138 |
| Strontium | **Inhibitory Concentration:** 10 mM Sr(II) | 139 |
| Vanadium | **Inhibitory Concentration:** 1.5 mM V(IV) | 140 |
| Zirconium | **Inhibitory Concentration:** 80 mM Zr(IV) | 141 |
| Aluminium | **Inhibitory Concentration:** 75 mM Al(III) | 142 |

## APPENDIX I. LEACHING STUDY WITH COMMERCIAL MARS REGOLITH SIMULANTS

### INTRODUCTION

Many published studies investigating chemical habitability in an aqueous Martian environment utilize extracts from leached regolith simulants as an available ISRU media component. Both the choice of regolith simulant and variations to leaching conditions will substantially impact the soluble inventory of chemicals. To expand our understanding of aqueous geochemistry in regolith extracts and inform future modifications to the Defined Mars Media (DMM) simulant presented in the main text of this report, we did a leaching study with different commercially available simulants across a range of conditions. The goal of this study was to identify which factors should be considered to reduce variation in soluble chemicals in leaching extracts.

### LEACHING STUDY

We conducted a study of existing standard simulants, leaching them in a range of regolith concentrations and pHs over different time periods (Appendix Table 1). We used three simulants that are commonly used in astrobiology research; JEZ-1 and MGS-1 from Space Resource Technologies and MMS-2 from The Martian Garden. We also looked into potential batch-to-batch variation by comparing the current with older batch preparations. Leachates were prepared using the same protocol depending on the variable being tested. Leachates were sent for ionic analysis via ion chromatography for major soluble anions (Cl^−^, NO_3_^−^, PO^3−^, SO_4_^2−^) and elemental analysis via inductively coupled plasma mass spectrometry (ICP-MS) for individual elements. Throughout this study, we classify the measured elements and ions into to three groups:

- The major soluble cations and anions measured *in situ* on Mars or in solubility studies: **Na, K, Ca, Mg, Cl**^**−**^, **NO**_**3**_^**−**^, **PO**^**3−**^, **SO**_**4**_^**2−**^
- Biologically essential trace elements: **Mn, Cu, W, B, Fe, Zn, Ni, Mo, Co, Se**
- Other trace elements incorporated into **DMM** (see main text): **Li, Al, Cr, Ba, V, Sr, Zr, Ag**

Note that individual elements are reported as total element concentration as ICP-MS can not determine charge state. The following elements were also measured via ICP-MS but not included in the figures or incorporated into DMM: Be, As, Te, Cs, Ta, Pb, Th, U, Si, Ti, Cd, Sb, Sn

### LEACHING SOLUTION PH

Studies that use regolith extracts for microbial growth often leach in a solution other than neutral water, such as a prepared growth media or acidic conditions to mimic biomining. To study the impact of the leaching solution (leachant) pH on the concentration of soluble species, we leached three regolith simulants in a range of pHs, using concentrated or dilute HCI and NaOH for acidic and basic conditions respectively, while keeping the input concentration constant. In acidic leaching conditions, the soluble concentration of most metal elements increases drastically, exhibiting classical dissolution behavior^143-144.^ Total calcium (Ca) and magnesium (Mg) increase in soluble concentration for all simulants in more acidic leaching conditions (Appendix Figure 1A). Similarly, the total soluble concentration of essential trace metals like manganese (Mn), iron (Fe), and cobalt (Co) increase by one to two orders of magnitude in acidic leaching conditions, depending on the simulant (Appendix Figure 1B). The MMS-2 simulant seemingly has less leachable total iron as the soluble iron concentration goes from ~0.011 mM at pH 7 to 0.2 mM at pH 1 (an 18-fold increase) while MGS-l and JEZ-1 increase from ~0.002 mM at pH 7 to 4.7 mM and 2.4 mM at pH 1 (a 2,350-fold and 1,200-fold increase) respectively. This behavior continues for the other trace elements (Appendix Figure 1C). For example, total chromium (Cr), aluminum (Al), and barium (Ba) concentrations increase in acidic conditions. In basic conditions, the total soluble concentration of metals like magnesium (Mg) and manganese (Mn) decrease significantly, and Mg is undetectable in MGS-1 and MMS-2 at pH 12, likely due to formation of insoluble hydroxides^145^. Meanwhile, oxophilic metals, like tungsten (W), increase in concentration in basic conditions^146^. There is a less drastic change for soluble (oxy)anions across different pHs (Appendix Figures 1A). Soluble sulfate remains between 3 - 10 mM at all pHs for all three simulants, except for MGS-1 leached at pH 1, which decreases to 0.95 mM. Soluble nitrate remains between 7.1 x 10^−2^ − 1.3 x 10^−1^ mM for MGS-1 and JEZ-1 simulants and between 1.98 x 10-2 - 2.5 x10^−2^ mM for MMS^−2^. In extremely acidic conditions (pH l), nitrate is not measured at all in MGS-1 and JEZ simulant extracts.

**Appendix Table 1.** Variables And Parameters Tested In Leaching Study.

| Variable | Parameters Tested |
| --- | --- |
| Simulant | MMS-2, MGS-1, JEZ-1 |
| Input regolith concentration (i.e. regolith:water ratio in g/L) | 1, 10, 40, 100, 200, 400, 800 |
| Leaching solution pH (i.e. leachant) | 1, 5, 7, 9, 12 |
| Leaching Time (days) | 1, 2, 5 |

**Appendix Figure 1.**
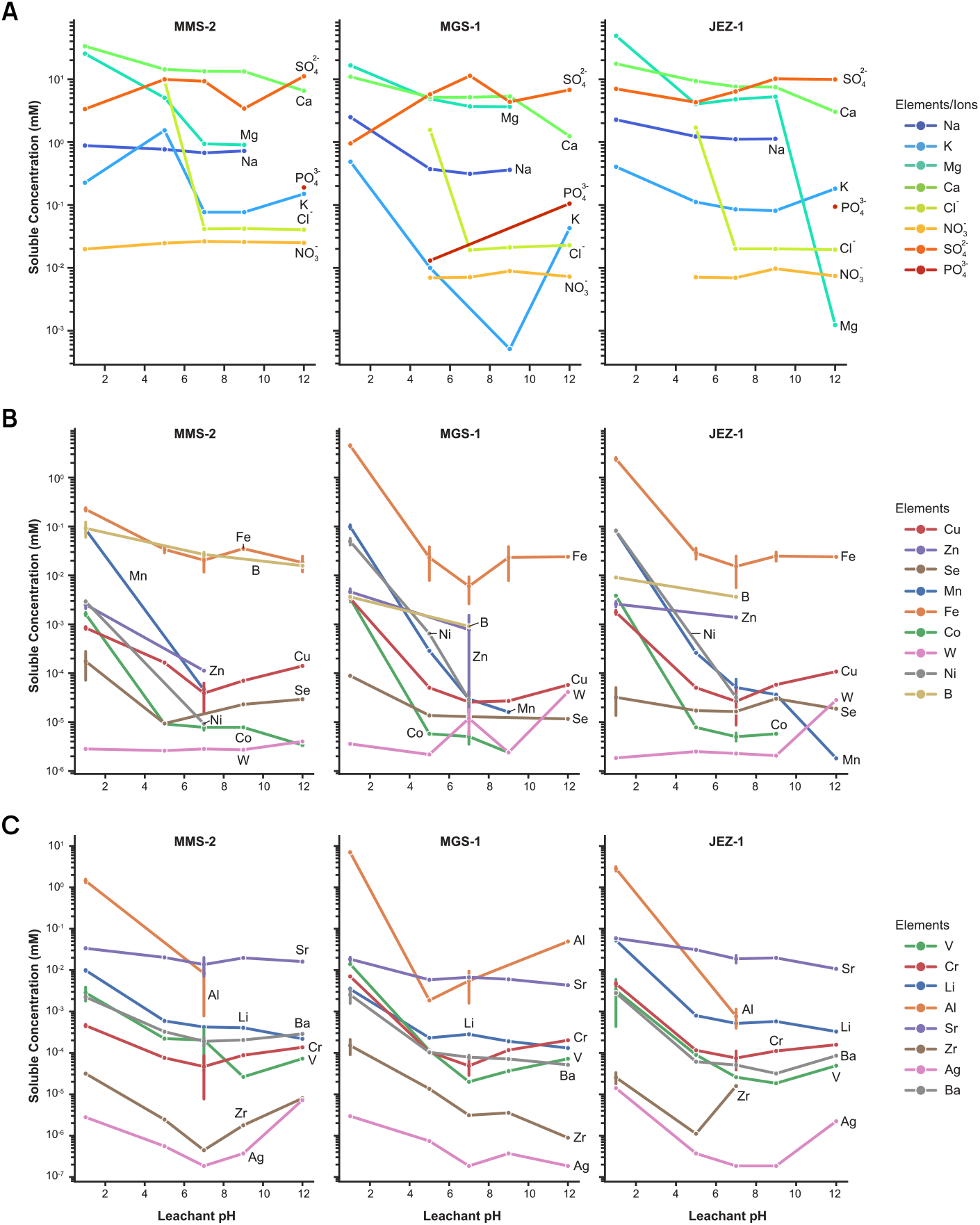
Impact of leaching solution pH on total soluble concentrations in extract. Concentration (in millimolar) of soluble species in leaching extracts for MMS-2. MGS-1, and JEZ-1leached at pH l. 5, 7. 9, and 12. Each dot is an average of 1-3 measurements depending on the sample, with error bars representing standard deviation where applicable. Colors represent different soluble species. Values for the concentration of total Na at pH 12 and Cl^−^ at pH 1 are removed as concentrated solutions of NaOH and HCl were used for leaching. A. Major soluble cations and anions B. Biologically essential trace elements C. Other trace elements For *all Figures, Ion Chromatography* is *used for* anion *(Cl*^−^, *NO*_*3*_^−^, Po_4_^−^,So_4_ ^2−^)*concentrations and ICP-MS* is *used for single element concentrations*.

### REGOLITH CONCENTRATION (I.E. REGOLITH:WATER RATIO)

We leached the simulants at a range of regolith to water ratios to measure how the concentration of soluble species changes with the amount of input regolith. Our baseline ratio is borrowed from WCL (40 g/L)^23^ and we tested up to 800 g/L regolith in water. In general, the total soluble concentration of elements increased with the ratio of regolith:water. In other words, when more regolith is added, more soluble elements are released and available. We expect at higher ratios that some elements may reach a solubility limit. For example, the concentration of total soluble calcium (Ca) in extracts of MGS-1 and MMS-2 leachates does not increase after 100 g/L as it plateaus at ~10-20 mM (**Appendix Figure 2A**). We see a similar pattern with magnesium (Mg), but only in MMS-2 extracts, where the total soluble concentration plateaus at ~1.5 mM after 100 g/L.

An expected shortcoming of these physical simulants is an overall low abundance of soluble anions like nitrate. However, we saw that the rate of nitrate scaled differently between the simulants. In MGS-1 extracts, nitrate levels look similar between 40 g/L to 400 g/L regolith concentration, at ~0.01 mM, while in MMS-2 extracts nitrate levels continue to increase, from 0.026 mM at 40 g/L to 0.12 mM at 200 g/L regolith concentration. Most soluble elements at trace levels also increase with increasing regolith concentrations. Some essential trace elements, such as Mo, Mn, Co, and Ni are only detected at input regolith concentrations at or above 40 g/L, likely indicating low total soluble abundance (Appendix Figure 2B). Many of the other trace elements, like strontium (Sr) and lithium (Li), also increase at a constant rate with increasing input amounts of regolith (Appendix Figure 2C). Similarly to calcium, the total soluble concentration of other trace elements decrease at the highest input regolith concentrations, like chromium (Cr), iron (Fe) and aluminum (Al), potentially due to a solubility limit or a higher concentration of counter ions with which it is insoluble (Appendix Figure 2B, 2C).

**Appendix Figure 2.**
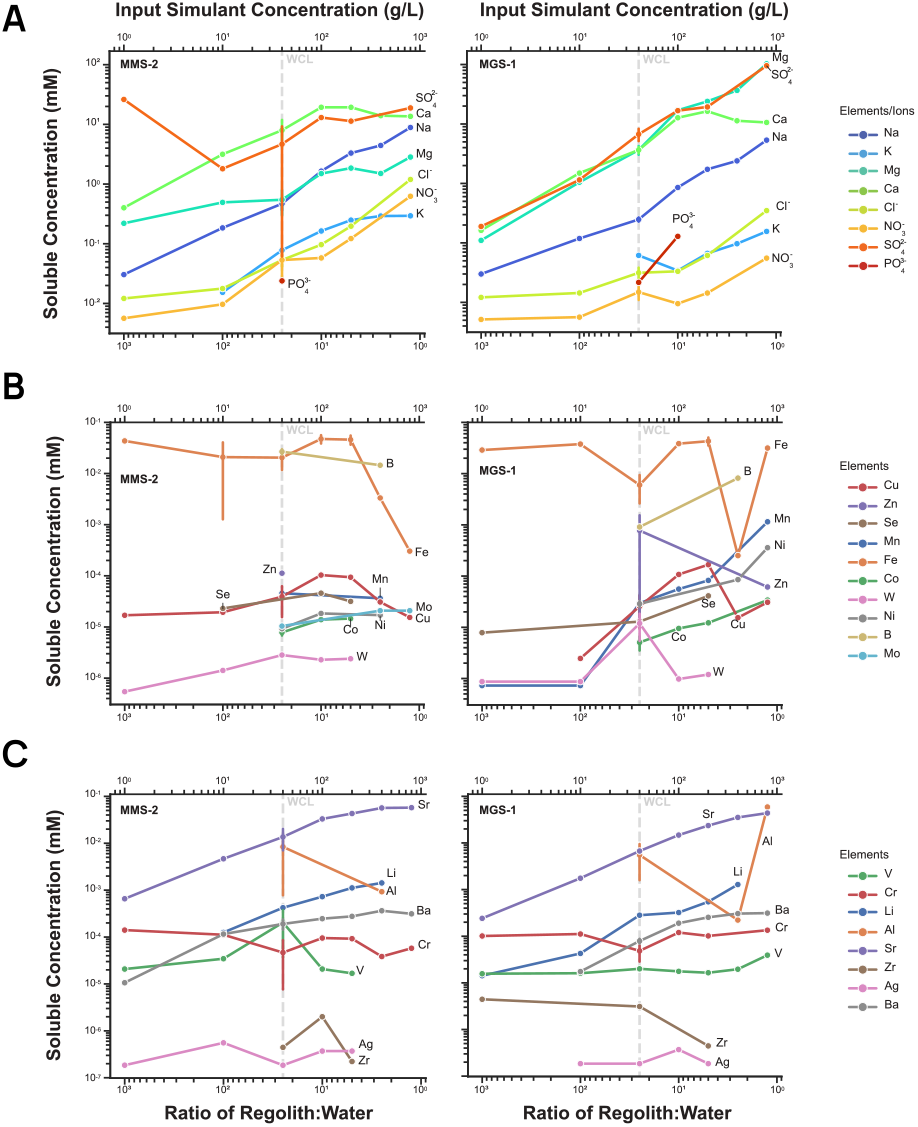
Impact of input regolith concentration (regolith:water ratio) on total soluble concentrations in extract. Concentration (in millimolar) of soluble species in extracts of MMS-2 and MGS-1leached at 1 g/L to 800 g/Linput regolith concentration (i.e. at regolith to water ratios of 1,000:l to l.25:l). Each dot is an average of 1 - 3 measurements depending on the sample, with error bars representing standard deviation where applicable. Colors represent different soluble species. Dotted line represents regolith:water ratio in the WCLexperiments(25:1 ratio of water to regolith, or 40 g/L regolith) A. Major soluble cations and anions B. Biologically essential trace elements C. Other trace elements *For all Figures, Ion Chromatography* is *used for anion (Cl*^−^, *NO*_*3*_^−^, Po_4_^−^,So_4_ ^2−^) *concentrations and ICP-MS* is *used for total element concentrations*.

### LENGTH OF LEACHING

Studies using regolith extracts as a basis for microbial growth often conduct leaching for anywhere between 12 - 72 hours and often do not describe the physical leaching method (ex. constant mixing, orbital shaking, etc.). To study the impact of length of leaching on the total soluble concentration of chemical species, we leached the MGS-l simulant for l, 2, and 5 days at constant orbital shaking at 40 g/L input regolith and in neutral water (pH 7). As regolith is leached longer, the total concentrations of most soluble species have minor changes and usually remain within the same order of magnitude (Appendix Figure 3). For example, the concentration of total soluble sodium (Na) increases by only ~6% from 1 day of leaching to 5 days of leaching (0.313 mM to 0.334 mM) (Appendix Figure 3A). The total soluble iron (Fe) concentration increases more drastically with longer leaching time, by ~50-60% (Appendix 3B). Other species like strontium (Sr) and lithium (Li) similarly increase by only ~1-3%. Some soluble elements decreased in total concentration, with zinc (Zn) decreasing from 3.05 x 10^-3^ mM at 1 day of leaching to 7.43 x 10^-5^ mM at 5 days of leaching (~97% decrease) (Appendix Figure 3C).

**Appendix Figure 3.**
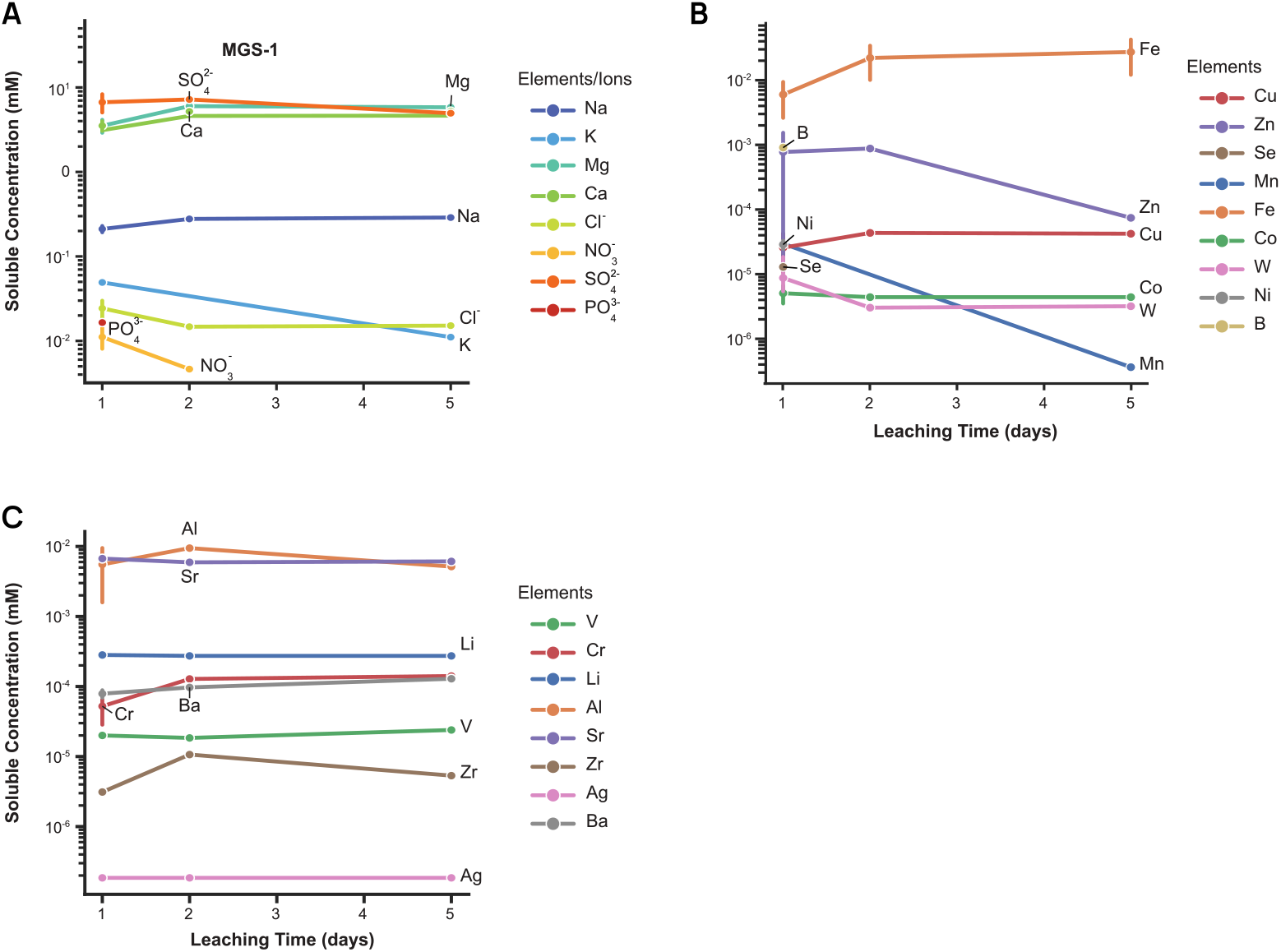
Impact of leaching time on total soluble concentrations in extract. Concentration (in millimolar) of soluble species in leaching extracts for MGS-1 leached at 40 g/L in pH 7 for 1, 2, or 5 days with continuous shaking. Each dot is an average of 1 - 3 measurements depending on the sample, with error bars representing standard deviation where applicable. Colors represent different soluble species. A. Major soluble cations and anions B. Biologically essential trace elements C. Other trace elements *For all Figures, Ion Chromatography* is *used* for anion *(Cl*^−^, *NO*_*3*_^−^, PO_4_^−^,SO_4_ ^2−^) concentrations *and ICP-MS* is *used* for *total element* concentrations.

### SIMULANT BATCH VARIATION

The commercially available simulants used in this study are all single or mixed source analogs. By virtue of being source analogs, new batches of simulants must be made after the initially collected sources run out. Simulant providers therefore update the batch or lot numbers depending on when the current batch’s source rocks were collected. There may inherently be some differences in source rocks collected at different times. We were interested in measuring the soluble species in different batches of different simulants to understand potential variation in studies using leachates of these simulants. We were able to source older batches of the three simulants we studied. All simulants were leached in neutral leaching conditions (40 g/L, pH 7 leachant) for 24 hours. Focusing on the major soluble species, there are slight but minimal differences in the total soluble concentration when comparing the current simulant batch to the older batches for JEZ-1 and MGS-1 (**Appendix Figure 4**). For example, the average concentration of total soluble sodium (Na) is ∼0.9 mM in leachates of the current batch of JEZ-1 and ∼1.5 mM in leachates of the older batch. Total soluble potassium (K) was less consistently measured in MGS-1 and MMS-2 simulant leachates compared to JEZ-1 simulant leachates. In MMS simulants, the largest differences in concentration between the current batch and the older batch is for total soluble magnesium (Mg), calcium (Ca) and sulfate (SO_4_^2−^). This difference may be explained by the differences in the simulant batches; the older batch tested was MMS-1 simulant, originally identified as a promising source based on its availability as whole rocks and dust and its hygroscopic behavior^33^, while the newer batch, MMS-2, includes added iron, silicon, magnesium, and calcium oxides to better mimic the composition of regolith^122^• There are also inherent differences between the simulants, again likely due to their mineralogical composition. In neutral leaching conditions, MGS-1 and JEZ-1 simulants have a higher concentration of total soluble magnesium and lower concentration of total soluble calcium compared to MMS-2. All three simulants are rich in soluble sulfate (SO_4_^2-^) but MMS-2 seems to have slightly more soluble chloride (Cl^-^) and nitrate (NO_3_^-^) compared to MGS-7 and JEZ-1.

**Appendix Figure 4.**
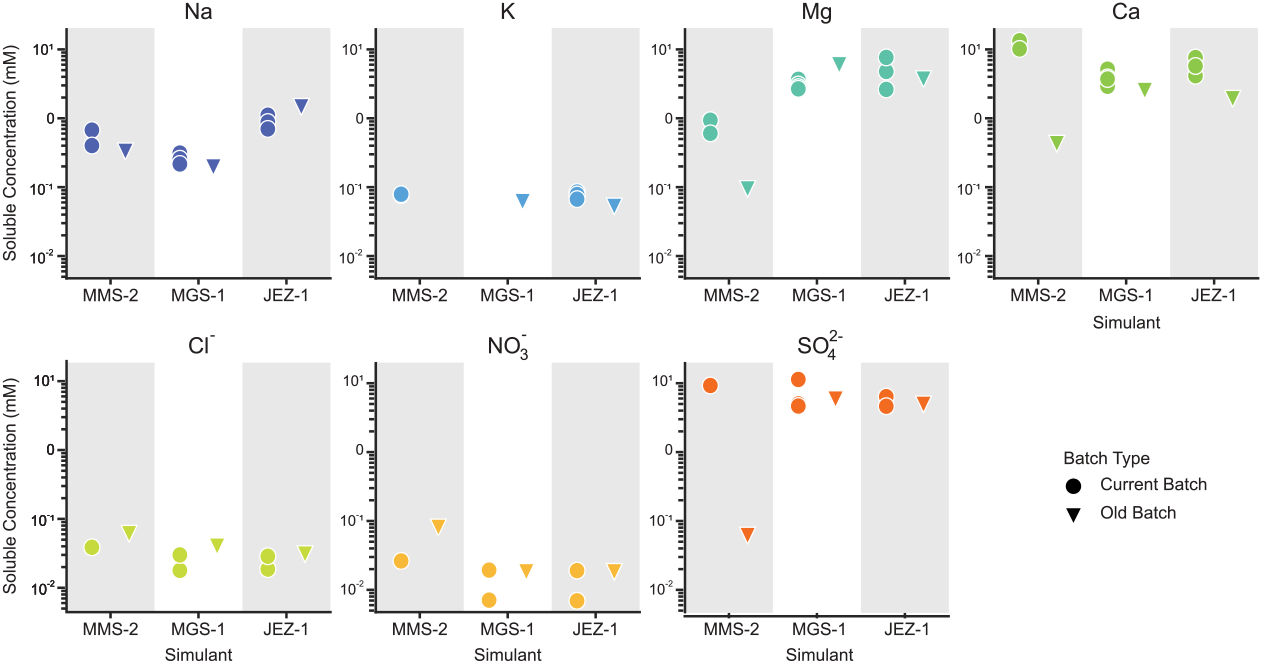
Impact of simulant batch on total soluble concentrations in extract Concentration (in millimolar) of soluble species in leaching extracts for MMS-2, MGS-1, and JEZ-1 leached at standard. A. conditions. See Supplementary Table 2 for batch number details. Each dot is a measurement for current batches and each triangle is a measurement for old batches. Colors represent different soluble species. *For all Figures, Ion Chromatography is used for anion (Cl^-^, NO*_*3*_, *PO*_*4*_ ^*3−*^, *SO*_*4*_ ^*2−*^*) concentrations and ICP-MS is used for total element concentrations*.

### CONCLUSION

We investigated how different leaching conditions influence the soluble concentration of different elements and ions across three regolith simulants. The largest influencing factor was the pH of the leaching solution (leachant), as many metals were highly enriched in more acidic leaching conditions and depleted in more basic conditions. Understanding regolith aqueous geochemistry in non-neutral pHs will help future plant habitability and mineral mining efforts. As the input regolith concentration increases in leaching, so too do the concentrations of most soluble metals. At some point, a solubility limit is reached, after which no more of that specific species will be soluble, or will even decrease at higher concentrations, as seen with calcium, magnesium, iron, copper, and sulfate. Studying the impact of regolith concentration can help identify ideal leaching conditions where highly soluble nutrients like nitrate can be obtained while still accurately modeling the concentration of stressors. like salts or perchlorate. The length of leaching or the specific simulant used had lesser effects on the total elemental concentrations. The differences between simulants is still essential to understand as different simulants are designed for specific purposes, such as a global simulant like MGS-1 or a site-specific simulant like JEZ-1. Other factors, like temperature and pressure, were not investigated and will also influence the dissolution behavior.

Understanding the available soluble molecules in regolith and their intrinsic solubility in relevant aqueous environments is essential for modeling an *in situ* feedstock. So far, leaching physical regolith simulants has been the only way to investigate this feedstock and the manner in which the leachates are prepared have a large impact on the chemical makeup of the resulting extracts. Studies such as this one will inform how to choose a leaching condition based on an experiment of interest and will dictate how aqueous simulants. like the DMM simulant presented in the main text, can be customized to match chosen leaching conditions. Using a consistent reference simulant will allow for more granular hypothesis testing based on available aqueous chemicals and standardize research across astrobiology laboratories.

All data used in this study is available as Appendix Dataset I: Ionic and Elemental Analysis Data.

## APPENDIX II. MODIFYING DMM FORMULATION TO REPRESENT DIFFERENT LEACHING CONDITIONS

In the main text, we report on Defined Mars Media (DMM), a defined salt mixture that acts as a simulant of soluble bio-available nutrients and stressors in leached Martian regolith **(Box 1**). We validated DMM’s composition up to a 20x concentrated version (i.e. 40 g/L to 800 g/L of input regolith concentration) and demonstrated its applicability in screening microbial chemical habitability. We also conducted a regolith simulant leaching study to inform how changing leaching conditions impacts the soluble makeup of the leachate**(Appendix I**). Identifying the impact of each condition on the leachate can help inform how DMM can be modified to represent those different conditions.

For example, at similar input regolith concentrations, the macronutrient concentration scaling is different between observed DMM and the simulants (Appendix Figure 5A). Concentrations of soluble ions at higher regolith concentrations do not scale linearly, while they do, as expected, in DMM. For example, the data suggests that the concentration of total soluble calcium in the tested physical simulants’ extracts does not increase when the input regolith concentration exceeds 100 g/L. Although the total soluble calcium concentration continues to rise in DMM, this value could be adjusted to the leaching limit when modifying DMM for higher regolith concentrations. A notable exception is total soluble magnesium and sulfate concentrations in MGS-1 extracts, which closely mirror the DMM concentrations.

In developing the micronutrient portion of DMM, we identified that many necessary trace elements are available at low but adequate soluble concentrations under neutral pH leaching conditions. When compared to nutrient requirement and growth inhibiting concentrations, total soluble concentrations of some micronutrients in acidic leaching conditions are no longer limiting and some are approaching inhibitory levels (**Appendix Figure 5B**). Under acidic leaching, required trace metals like Mn, Ni, and Cu are safely above the minimum required concentration for growth while Fe and Cr concentrations approach or exceed inhibitory concentrations. Soluble total molybdenum is no longer detected in acidic leaching conditions, but is likely to increase in alkaline conditions. These metals’ behaviors in acidic leaching are similar to nutrient availability behavior in acidic soil on Earth^147,148^•

DMM serves as a helpful baseline simulant from which future iterations, such as higher concentration and non-neutral leaching simulants, can be created. Modifying higher DMM concentrations to take solubility limits into account is as simple as lowering the input amount of calcium and sulfate while keeping the remaining ions in balance to maintain a pH of 7.7. Similarly, an acidic micronutrients solution can be prepared by formulating a new recipe based on the total soluble concentrations in acidic leaching conditions.

**Appendix Figure 5.**
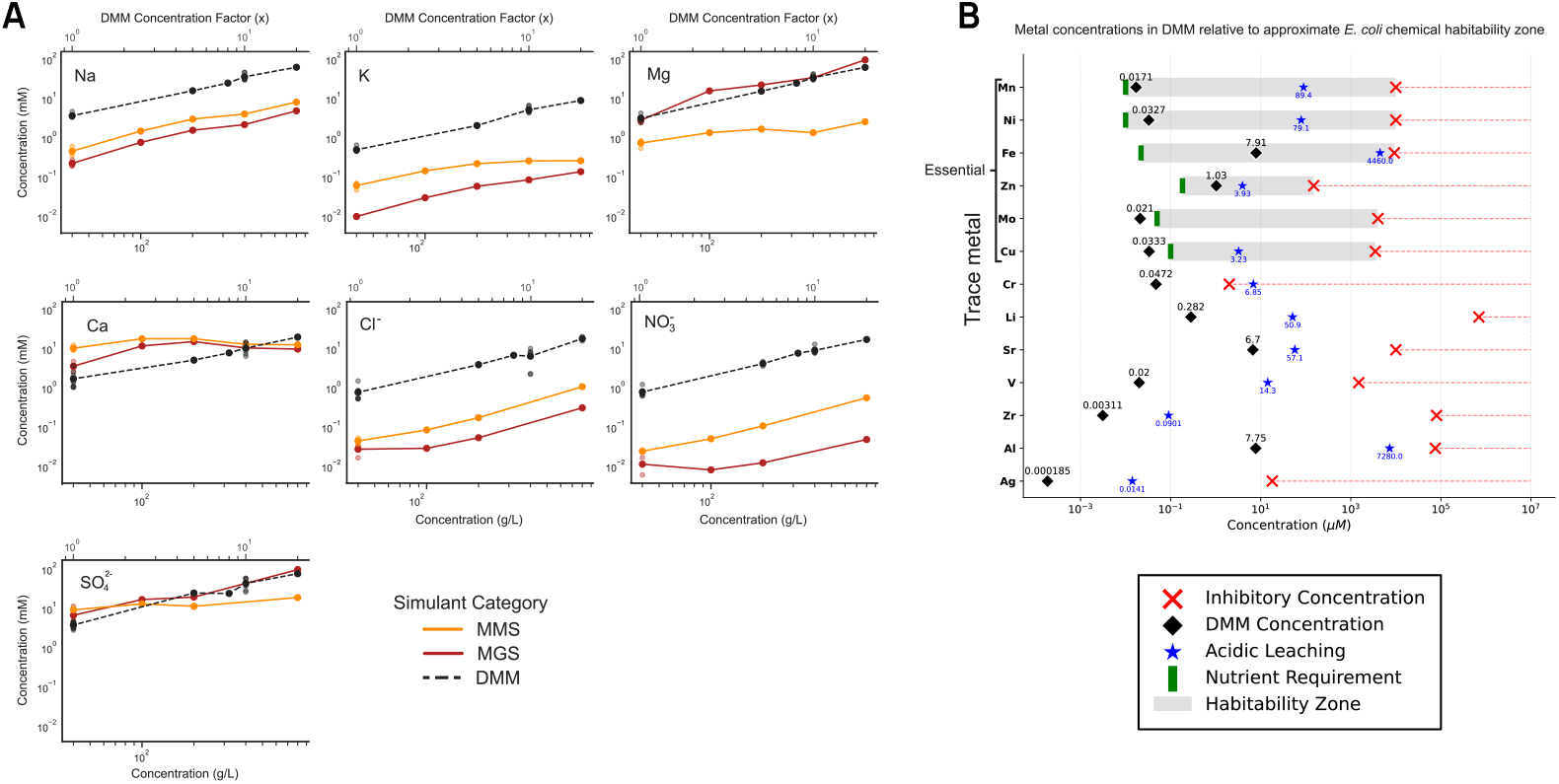
Comparison of total soluble concentrations from regolith simulation leaching to DMM values and biologically relevant concentrations. **A**. Concentration (in millimolar) of soluble species in extracts of MMS-2 (orange line) and MGS-1 (dark red line) leached at input regolith concentrations of 40 g/L to 800 g/L compared to the measured concentrations in DMM simulant at those same ratios (black dotted line), described by Concentration Factor of 1x to 20x. Each dot represents a single measurement and the line connects the averages for each concentration. Perchlorate is not shown because perchlorate was not detected in physical simulant leachates. Phosphate is not shown due to measurement difficulties with co-elution around the sulfate peak in ion chromatography. **B**. Comparison of total soluble concentration of trace elements in DMM (black diamond) and in acidic leaching of simulants (blue star) to nutrient requirement and inhibitory concentrations of those trace elements for *E. coli* **(see Supplementary Table 3)**. Molybdenum (Mo) was not detected in acidic leaching conditions. *For all Figures, Ion Chromatography is used for anion (Cl^-^, NO*_*3*_, *SO*_*4*_ ^*2−*^*) concentrations and ICP-MS is used for total element concentrations*.

